# Endomembranes promote chromosome missegregation by ensheathing misaligned chromosomes

**DOI:** 10.1101/2021.04.23.441091

**Authors:** Nuria Ferrandiz, Laura Downie, Georgina P. Starling, Stephen J. Royle

**Affiliations:** Centre for Mechanochemical Cell Biology, Warwick Medical School, Gibbet Hill Road, Coventry, CV4 7AL, UK; The Francis Crick Institute, London, NW1 1AT, UK

**Keywords:** chromosome missegregation, endoplasmic reticulum, mitotic spindle, chromosomal instability, cancer

## Abstract

Errors in mitosis that cause chromosome missegregation lead to aneuploidy and micronuclei formation which are associated with cancer. Accurate segregation requires the alignment of all chromosomes by the mitotic spindle at the metaphase plate, and any misalignment must be corrected before anaphase is triggered. The spindle is situated in a membrane-free “exclusion zone”, beyond this zone, endomembranes (endoplasmic reticulum, nuclear envelope and other organelles) are densely packed. We asked what happens to misaligned chromosomes that find themselves beyond the exclusion zone? Here we show that such chromosomes become ensheathed in multiple layers of endomembranes. Chromosome ensheathing delays mitosis and increases the frequency of chromosome missegregation. The micronuclei that form following missegregation have a disrupted nuclear envelope with internal endomembranes. We use an induced organelle relocalization strategy in live cells to show that clearance of endomembranes allows for the rescue of chromosomes that were destined for missegregation. Our findings indicate that endomembranes promote the missegregation of misaligned chromosomes that are outside the exclusion zone, and therefore constitute a risk factor for aneuploidy.

## Introduction

Accurate chromosome segregation during mitosis is essential to prevent aneuploidy, a cellular state of abnormal chromosome number (Duijf and Benezra, 2013). Errors in mitosis that lead to aneuploidy can occur via different mechanisms. These mechanisms include: mitotic spindle abnormalities (Ghadimi et al., 2000), incorrect kinetochore-microtubule attachments (Cimini et al., 2001), dysfunction of the spindle assembly checkpoint (Kalitsis et al., 2000), defects in cohesion (Daum et al., 2011) and failure of cytokinesis (Fujiwara et al., 2005). Some of these error mechanisms result in the missegregation of whole chromosomes, a process termed chromosomal instability (CIN). The majority of solid tumors are aneuploid, with higher rates of CIN and so understanding the mechanisms of chromosome missegregation is an important goal of cancer cell biology. In addition, chromosome missegregation is associated with micronuclei formation, which is linked to genomic rearrangements that may drive tumour progression (Crasta et al., 2012; Ly et al., 2017; Liu et al., 2018). While the mitotic spindle has logically been the focus of efforts to understand chromosome missegregation, there has been less attention on other features of mitotic cells such as intracellular membranes. In eukaryotic cells, entry into mitosis constitutes a large scale reorganization of intracellular membranes. The nuclear envelope (NE) breaks down while the endoplasmic reticulum (ER) and Golgi apparatus disperse to varying extents (Hepler and Wolniak, 1984; Warren, 1993). These organelle remnants – collectively termed “endomembranes” – are localized toward the cell periphery while the mitotic spindle itself is situated in an “exclusion zone” which is largely free of membranes and organelles (Bajer, 1957; Porter and Machado, 1960; Nixon et al., 2017). The endomembranes beyond the exclusion zone are densely packed although the details of their ultrastructure varies between cell lines (Puhka et al., 2007; Lu et al., 2009, 2011; Puhka et al., 2012; Champion et al., 2017). This arrangement means that, although mitosis is open in mammalian cells, the spindle operates within a partially closed system. Several lines of evidence suggest that endomembranes must be cleared from the exclusion zone in order for the mitotic spindle to function normally (Vedrenne et al., 2005; Schlaitz et al., 2013; Champion et al., 2019; Kumar et al., 2019; Merta et al., 2021). In addition, it is thought that this arrangement is required to concentrate factors needed for spindle formation (Schweizer et al., 2015). This study was prompted by a simple question: what happens to misaligned chromosomes that find themselves beyond the exclusion zoneã We show that such chromosomes become ensheathed in multiple layers of endomembranes. Chromosome ensheathing delays mitosis and increases the frequency of chromosome missegregation and subsequent micronuclei formation. Using an induced organelle relocalization strategy we demonstrate that clearance of endomembranes allows the rescue of chromosomes that were destined for missegregation. Our findings indicate that endomembranes are a risk factor for CIN if the misaligned chromosomes go beyond the exclusion zone boundary during mitosis.

## Results

### Misaligned chromosomes outside the exclusion zone are ensheathed in endomembranes

During mitosis, the spindle apparatus is situated in a membrane-free “exclusion zone”. Outside the exclusion zone, the ER and nuclear envelope – collectively called “endomembranes” – surround the mitotic spindle. We investigated the organization of endomembranes in mitotic cells using light and electron microscopy. First, we carried out live-cell imaging of mitotic RPE-1 cells that stably co-express GFP-Sec61β and Histone H3.2-mCherry, stained with SiR-tubulin to mark the ER, DNA and microtubules, respectively. These images revealed a mitotic spindle-sized exclusion zone from which GFP-Sec16β signal was absent (Figure 1A). Second, serial block face-scanning electron microscopy (SBF-SEM) of mitotic RPE-1 cells showed that the ellipsoid exclusion zone is largely devoid of endomembranes, including mitochondria and other organelles. Outside the exclusion zone, endomembranes are tightly packed and the border between these two regions is clearly delineated and could be segmented (Figure 1B).

**Figure 1.**
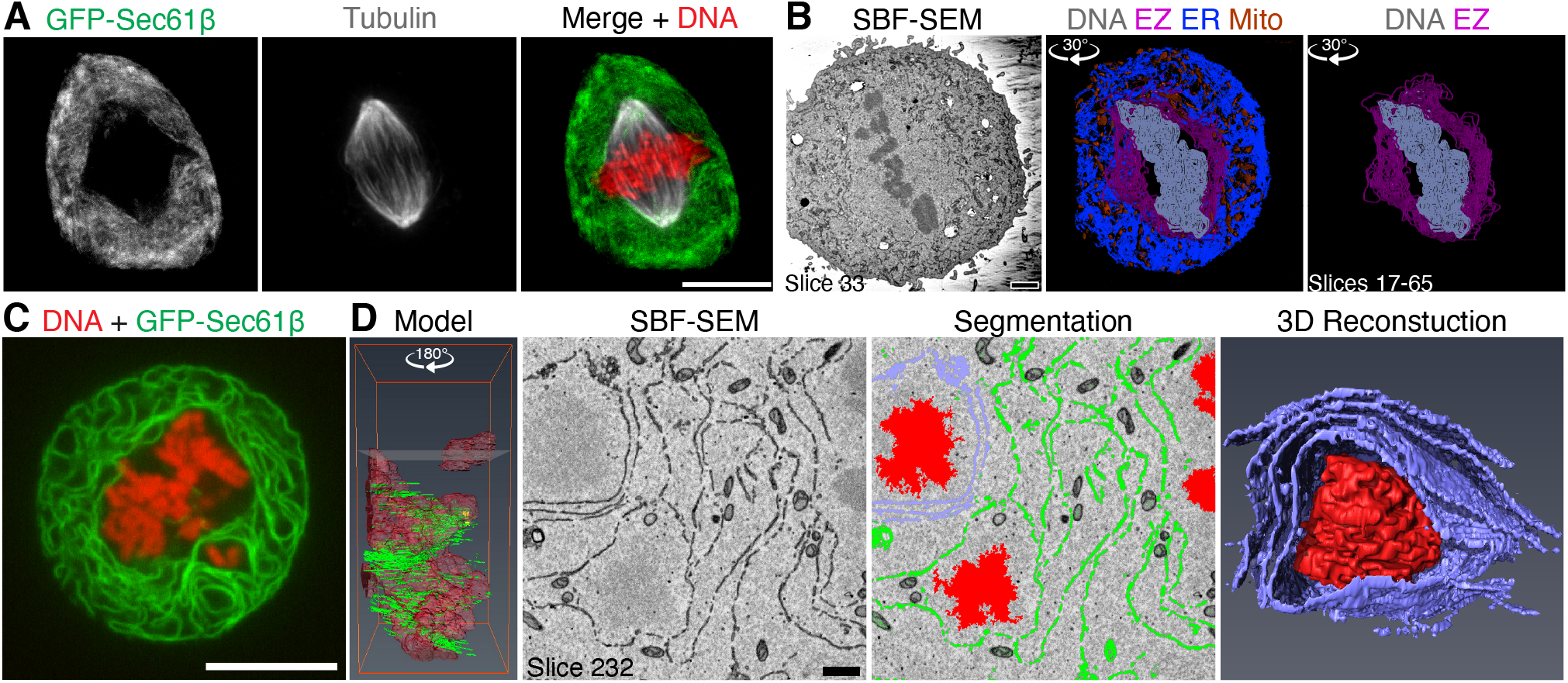
Misaligned chromosomes outside the exclusion zone are ensheathed in endomembranes. (**A**) Confocal image of a mitotic RPE-1 cell stably co-expressing GFP-Sec61β (green) Histone H3.2-mCherry (DNA, red) and stained with SiR-Tubulin (gray). Scale bar, 10 µm. (**B**) Serial block face-scanning electron microscopy (SBF-SEM) imaging of mitotic cells and subsequent segmentation reveals the endomembranes (ER, blue) and mitochondria (Mito, orange) are beyond the exclusion zone boundary (EZ, pink), with the chromosomes (DNA, gray) within. Angle of rotation about Y axis is shown. Scale bar, 2 µm. (**C**) Confocal image of an untreated HeLa cell co-expressing Histone H2B-mCherry and GFP-Sec61β with a spontaneously occurring ensheathed chromosome. (**D**) SBF-SEM imaging of an untreated HeLa cell with a spontaneously occurring ensheathed chromosome. Model shows the position of two ensheathed chromosomes (red) away from the metaphase plate, height of slice 232 is indicated. Segmentation shows endomembranes (green and lilac surrounding the upper chromosome), rendered in 3D (Reconstruction). Scale bar, 1 µm. See Supplementary Videos SV1 and SV2.

Misaligned chromosomes are those that fail to attach, or lose their attachment to the mitotic spindle. What happens to misaligned chromosomes that find themselves amongst the endomembranes beyond the exclusion zoneã HeLa cells have high rates of chromosome misalignment and live cell imaging showed that misaligned chromosomes could be situated beyond the exclusion zone (Figure 1C). Reconstruction of SBF-SEM data from HeLa cells showed that three to four layers of endomembranes “ensheath” the chromosomes beyond the exclusion zone (Figure 1D and Supplementary Video SV1 and SV2). We use the term “ensheathed” to describe how these chromosomes are surrounded by endomembranes but not fully enclosed in any one layer as though in a vesicle.

In order to study chromosome ensheathing in diploid cell lines, we needed to artificially increase the frequency of misaligned chromosomes in mitosis. Our main model was RPE-1 cells pre-treated with 150 nM GSK923295, a CENP-E inhibitor (Wood et al., 2010), before washing out the drug for 1 h (Figure 2A). In parallel, we also used a system of targeted Y-chromosome spindle detachment in DLD-1 cells (Ly et al., 2017) (Figure S1). Using live-cell imaging in both cell types, we observed that misaligned chromosomes beyond the exclusion zone are submerged in endomembranes (Figure 2B and S1D). Next, we used an image analysis method to determine the location of kinetochores in 3D space and map these positions relative to the exclusion zone boundary (see Methods, Figure 2D and S1E). Kinetochores of chromosomes that were not aligned at the metaphase plate therefore fell into two categories: those that were surrounded by GFP-Sec61β signal, termed ‘ensheathed’ and those that were not, termed ‘free’ (Figure 2B,D). Spatial analysis revealed that the kinetochores of ensheathed chromosomes were beyond the exclusion zone, whereas kinetochores of free chromosomes lay at the boundary in RPE-1 cells (Figure 2E). In DLD-1 cells the distinction was even more clear, with the kinetochores of free chromosomes positioned inside the exclusion zone S1F). The exclusion zone therefore approximately defines chromosome misalignment, with those chromosomes beyond the exclusion zone likely to be ensheathed by endomembranes. However, imaging GFP-Sec61β was required to verify that a chromosome was fully ensheathed.

**Figure 2.**
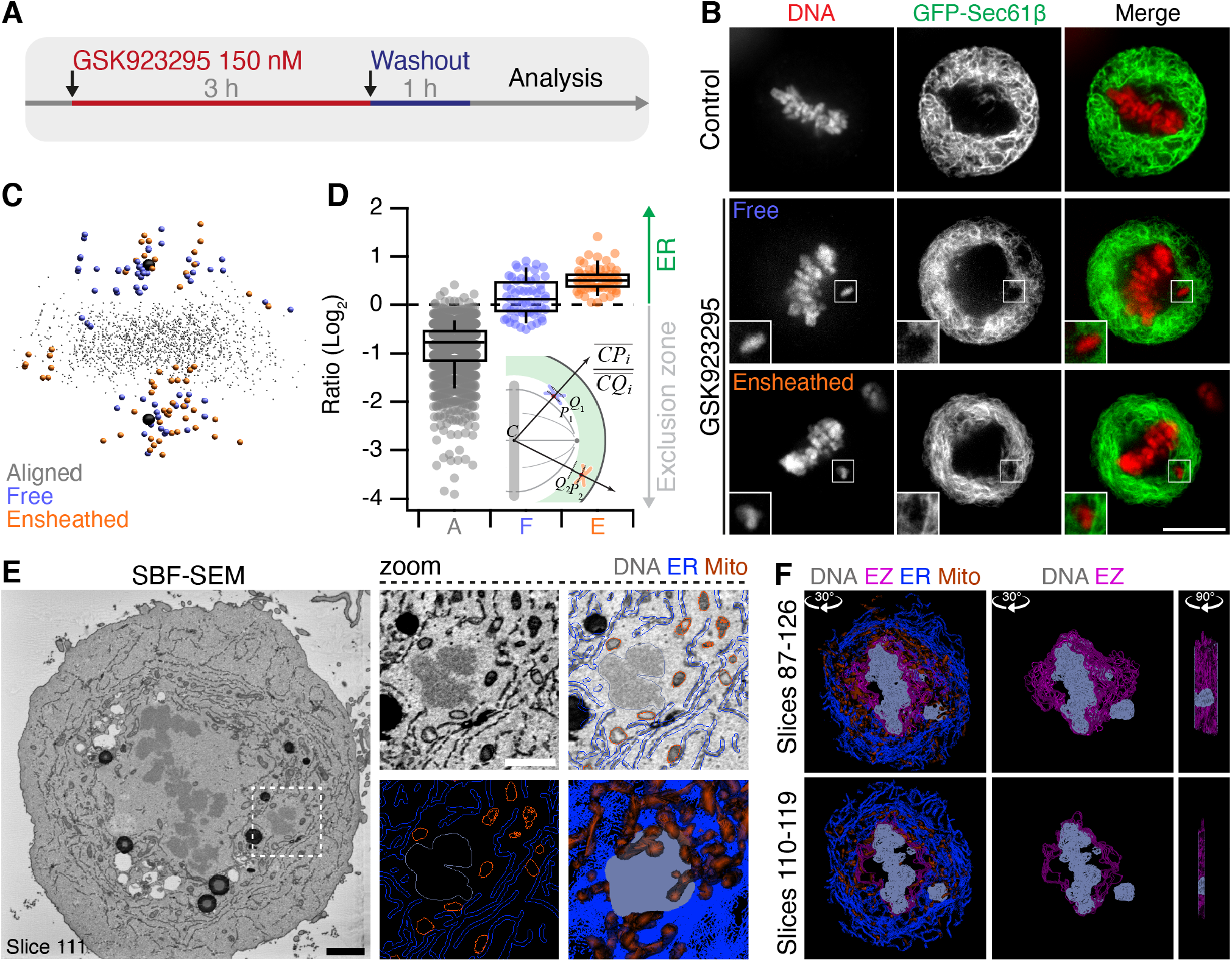
Induction of misaligned chromosomes in stably diploid RPE1 cells by pre-treatment with a CENP-E inhibitor. (**A**) Polar, misaligned chromosomes can be induced by treatment with CENP-E inhibitor GSK923295 (150 nM, 3 h) and subsequent washout (1 h). (**B**) Confocal micrographs to show that these misaligned chromosomes (SiR-DNA, red) are either outside the exclusion zone delineated by GFP-Sec61β (green), termed *Ensheathed* or at the boundary and inside the exclusion zone, termed *Free*. Scale bar, 10 µm. (**C**) Spatially averaged 3D view of all CENP-C-positive kinetochores in the dataset (see Methods). Small gray points represent kinetochores at the metaphase plate. Colored points represent misaligned chromosomes that were ensheathed (orange) and those that were not (Free, blue). Spindle poles are shown in black. (**D**) Box plot to show the relative position of each kinetochore relative to the exclusion zone boundary. Chromosome misalignment was induced pre-treatment with GSK923295 (150 nM). Ratio of kinetochores within the exclusion zone are *<* 0 and those within the ER are *>* 0 on a log2 scale. Dots represent kinetochore ratios from 31 RPE-1 cells at metaphase. Boxes show IQR, bar represents the median and whiskers show 9th and 91st percentiles. Inset: Schematic diagram to show how the position of kinetochores relative to the exclusion zone boundary was calculated. C is the centroid of aligned kinetochores, P is a kinetochore and Q is the point along the 3D path (CP) that intersects the exclusion zone boundary. The ratio of CP to CQ is taken for each kinetochore (aligned kinetochores, gray; free, blue; and ensheathed, orange). (**E**) Single SBF-SEM image showing an ensheathed chromosome. Boxed region is shown expanded and modeled (zoom). Single slice and a 3D model (bottom right) of slices 87-126 are shown. Scale bar, 2 µm (black) and 500 nm (white). (**F**) Modeled substacks from SBF-SEM images showing a chromosome outside the exclusion zone, ensheathed in ER. Slices shown and angles and axes of rotation are indicated (See Supplementary Videos SV3).

We again used SBF-SEM to observe how chromosomes beyond the exclusion zone interact with endomembranes in RPE-1 cells. Cells observed by fluorescence microscopy to have at least one ensheathed chromosome were selected for 3D EM analysis (Figure 2E). Segmentation of these datasets confirmed that the chromosome was fully beyond the exclusion zone boundary (Figure 2F and Supplementary Video SV3) and was ensheathed in several layers of endomembranes (Figure 2E).

The observation of ensheathed chromosomes raised immediate questions about their fate and whether ensheathing leads to aberrant mitosis.

### Ensheathed chromosomes delay mitotic progression

To determine the impact of ensheathed chromosomes on cell division, we first analyzed mitotic progression in RPE-1 cells stably expressing GFP-Sec61β with induction of ensheathed chromosomes using GSK923295 pre-treatment. Cells that had at least one ensheathed chromosome showed prolonged mitosis; median nuclear envelope breakdown (NEB)-to-anaphase timing of 66 min compared with 27 min in GSK923295 pre-treated cells where all chromosomes were aligned. The time to align the majority of chromosomes (NEB- to-metaphase) was delayed for cells with either a free or an ensheathed chromosome, but cells with an ensheathed chromosome had an additional delay to progress to anaphase (Figure 3A). Given these delays, we next confirmed that the spindle assembly checkpoint was active in these cells. The amount of Mad2 and Bub1 detected by immunofluorescence at CENP-C positive kinetochores of free or ensheathed chromosomes was similar, and four-fold higher than at kinetochores of aligned chromosomes (Figure 3B,C and S2A,B, for DLD-1 cells). Using live cell imaging, we found that GFP-Mad2 was recruited to kinetochores of ensheathed chromosomes (Figure 3D,E). Semi-automated 4D tracking of chromosomes allowed us to monitor their GFP-Mad2 status over time, relative to anaphase onset. These data revealed that GFP-Mad2 is lost from ensheathed chromosomes with similar kinetics to the signals at misaligned chromosomes that successfully congress to the metaphase plate (Figure 3E).

**Figure 3.**
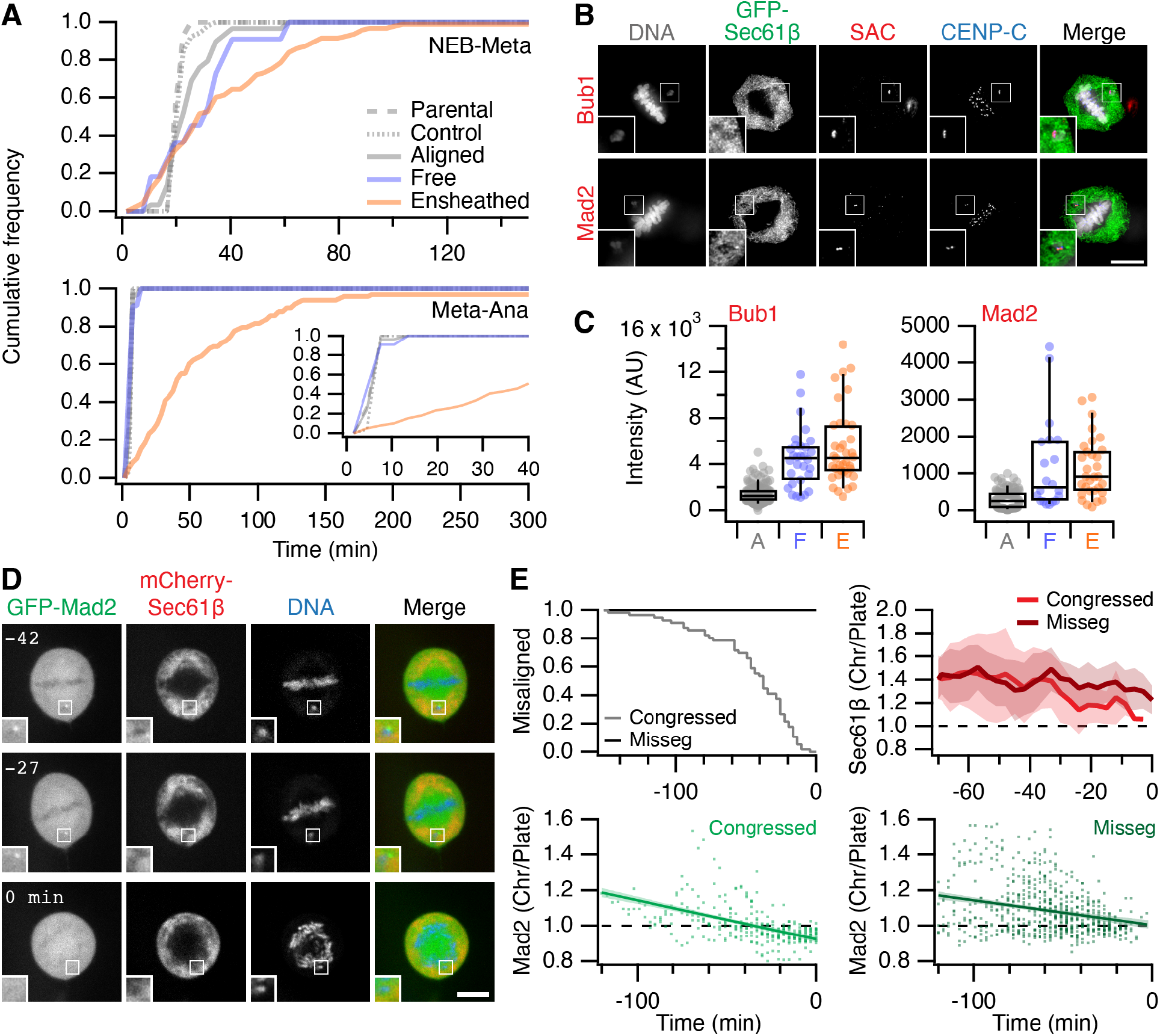
Impact of ensheathed chromosomes on cell division. (**A**) Mitotic timing of RPE-1 cells. Cumulative frequencies for nuclear envelope breakdown to metaphase (NEB-Meta) and metaphase to anaphase (Meta-Ana) are shown. RPE-1 stably expressing GFP-Sec61β were treated with 150 nM GSK923295 for 3 h before washout. Three classes of metaphase were seen: all chromosomes aligned (Aligned, *n* = 29), cells with one or more free chromosome (Free, *n* = 11), and cells with one or more ensheathed chromosome (Ensheathed, *n* = 107). Timing of untreated parental (Parental, *n* = 69) and stable RPE-1 (Control, *n* = 52) cells is also shown. Inset in Meta-Ana shows same data on an expanded timescale. Comparison of NEB-Meta and Meta-Ana timing distributions for Ensheathed vs Control, p = 1.9 *×* 10^−57^ and 7.8 *×* 10^−23^, Kolmogorov-Smirnov test. (**B**) Micrographs of immunofluorescence experiments to detect Bub1 or Mad2 (SAC, red) at kinetochores (CENP-C, blue) in cells stably expressing GFP-Sec61β (green), DAPI-stained DNA is shown in gray. Scale bar, 10 µm. (**C**) Quantification of Bub1 and Mad2 immunofluorescence at kinetochores marked by CENP-C. Ensheathed chromosomes were classified using the GFP-Sec61β signal. Dots represent cells, boxes show IQR, bar represents the median and whiskers show 9th and 91st percentiles. (**D**) Stills from live cell imaging experiments to track Mad2 levels at kinetochores of ensheathed chromosomes. A GSK923295-pre-treated RPE-1 cell is shown, stable co-expressing GFP-Mad2 (green) and mCherry-Sec61β (red), DNA is stained using SiR-DNA (blue). Time relative to anaphase is shown in minutes. Insets show 2X zoom of the indicated ROI. Scale bar, 10 µm. (**E**) Quantification of live Mad2 imaging experiments. Kaplan-Meier plot to show congression times of the last misaligned chromosome to align. Measurement of mCherry-Sec61β (mean ± s.d.) and GFP-Mad2 is shown for the misaligned that congressed and those that were missegregated (misseg). A linear regression fit with 95 % confidence intervals is shown for GFP-Mad2. All plots are shown in time in minutes relative to anaphase onset. Total cells with misaligned chromosomes, *n* = 72; cells where all chromosomes congressed, *n* = 56; and where there was missegregation (misseg), *n* = 16.

The failure of ensheathed chromosomes to congress is likely due to a lack of microtubule attachment, suggesting that endomembranes inhibit chromosome-microtubule interactions. We confirmed that ensheathed chromosomes have no stable end-on kinetochore-microtubule attachments by detecting colocalization of kinastrin, a marker for stable end-on attachment (Dunsch et al., 2011), with kinetochores of aligned and misaligned chromosomes (Figure S4A-C). Live cell imaging of RPE-1 cells stably co-expressing Histone H3.2-mCherry and GFP-Sec61β, stained with SiR-Tubulin showed that ensheathed chromosomes that failed to congress had no detectable microtubule contacts; whereas free chromosomes that had microtubule contacts, could be rescued and aligned at the metaphase plate, albeit after a delay (Figure S4D,E).

These results suggest that ensheathed chromosomes hinder mitotic progression in a spindle assembly checkpoint-dependent manner. Lack of microtubule contact is sensed by the spindle assembly checkpoint, but ultimately, the checkpoint is extinguished in the absence of congression after a long delay. The cells then proceed to anaphase resulting in missegregation of the ensheathed chromosome.

### Ensheathed chromosomes promote micronuclei formation

To understand the fate of cells with an ensheathed chromosome, we next examined mitosis in control or GSK923295-pre-treated RPE-1 cells stably expressing GFP-Sec61β using live-cell spinning disc microscopy (Figure 4A). In cells with an ensheathed chromosome, we observed the long delay in mitosis relative to control cells, and that mitosis was often resolved by missegregation and formation of a micronucleus (Figure 4A and S2C for DLD-1 cells). These experiments suggested that ensheathed chromosomes are potentially a precursor to micronuclei. We therefore followed the fate of mitotic cells by long-term live-cell imaging in order to understand the likelihood of mitotic outcomes. Our sample of cells pre-treated with GSK923295 included the three metaphase classes: aligned (25.8 %), free (5.4 %), and ensheathed (65.6 %). The most frequent fate of cells with an ensheathed chromosome was micronucleus formation (39 %). Of the 47 cells that formed a micronucleus after division in the dataset, 46 were from the ensheathed class (Figure 4B). This promotion of micronucleus formation was significant in cells with an ensheathed chromosome compared to free (p = 1.3 × 10^−3^, Fisher’s exact test). A smaller proportion of cells with an ensheathed chromosome exited mitosis normally, albeit with a delay (34 %), with the remainder showing other defects or death (20 % or 8 %). Cells pre-treated with GSK923295, that had aligned all their chromosomes, had similar fates to parental and control cells (Figure 4B and Supplementary Video SV5 and SV6). These fate-mapping experiments suggest that ensheathing of chromosomes by endomembranes promotes the formation of micronuclei.

**Figure 4.**
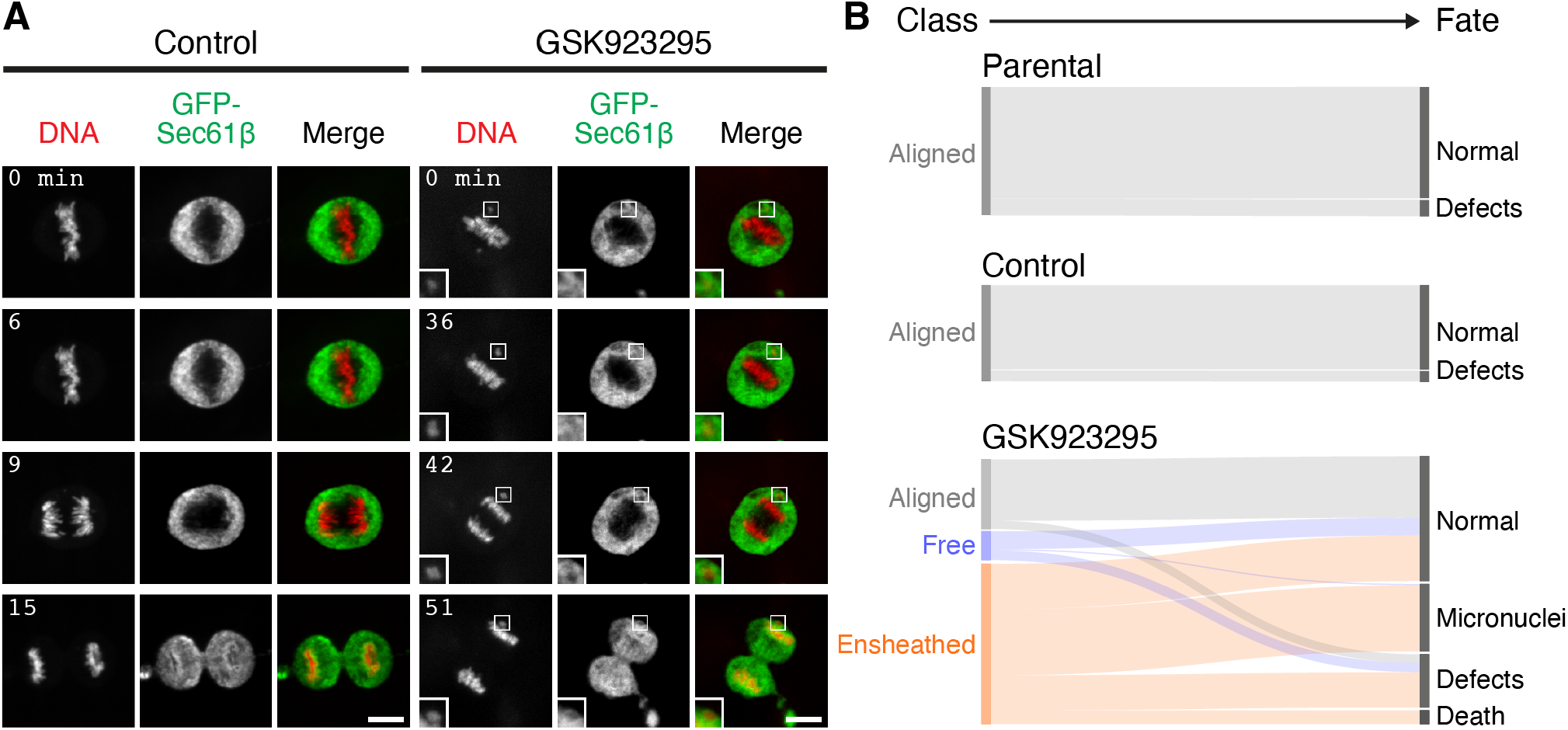
Ensheathed chromosomes promote micronuclei formation. (**A**) Stills from live cell imaging experiments to track the fate of ensheathed chromosomes. A control or GSK923295-pre-treated GFP-Sec61β RPE-1 cell is shown, DNA is stained using SiR-DNA (red). Scale bar, 10 µm. Shown in Supplementary Videos SV5 and SV6. (**B**) Sankey diagram to show the fate (right) of cells in each of the three metaphase classes (left). Fates include normal division, micronuclei formation, death and other defects (lagging chromosome, cytokinesis failure). Note that the fate of cells (and not chromosomes) is tracked. A cell with three misaligned chromosomes, only one of which is ensheathed is classified as Ensheathed. Parental RPE-1 cells (Parental, *n* = 92) and untreated RPE-1 stably expressing GFP-Sec61β (Control, *n* = 69) are from two and three independent overnight experiments, respectively. Fates of GSK923295-pre-treated GFP-Sec61β cells (*n* = 186) were compiled from seven experiments. Fates of individual chromosomes are shown in Supplementary Figure S3B)

### Micronuclei formed from ensheathed chromosomes have a disrupted nuclear envelope

Micronuclei can undergo a collapse of their nuclear envelope which manifests as ER tubules invading the micronuclear space (Hatch et al., 2013). We therefore asked if micronuclei that formed from ensheathed chromosomes were similarly defective. Using confocal imaging of RPE-1 cells stably co-expressing GFP-Sec61β and either mCherry-BAF or LBR-mCherry that were fixed 8 h after washout of GSK923295 to examine micronucleus integrity, we found the majority of micronuclei have ER inside the micronucleus (Figure 5). The fluorescence of GFP-Sec61β was higher at the micronucleus when compared with the main nucleus (Figure 5B). Moreover, the levels of either mCherry-BAF or LBR-mCherry were correlated with GFP-Sec61β. To confirm that these micronuclei had disrupted nuclear envelopes, we stained for H3K27Ac, a modification to Histone H3 that is removed by exposure to the cytoplasm (Mammel et al., 2021). Intact micronuclei had similar H3K27Ac signals to the corresponding main nucleus, whereas in micronuclei that were disrupted, the signal was lost (Figure 5A). The ratio of H3K27Ac signal at the micronucleus compared with the main nucleus was anticorrelated with the ratios of GFP-Sec61β, mCherry-BAF, and LBR-mCherry (Figure 5B). Since the majority of micronuclei formed following pre-treatment of RPE-1 cells with GSK923295 are derived from ensheathed chromosomes (Figure 4B), these data suggest that the ensheathing process may contribute to the formation of defective micronuclear envelope.

**Figure 5.**
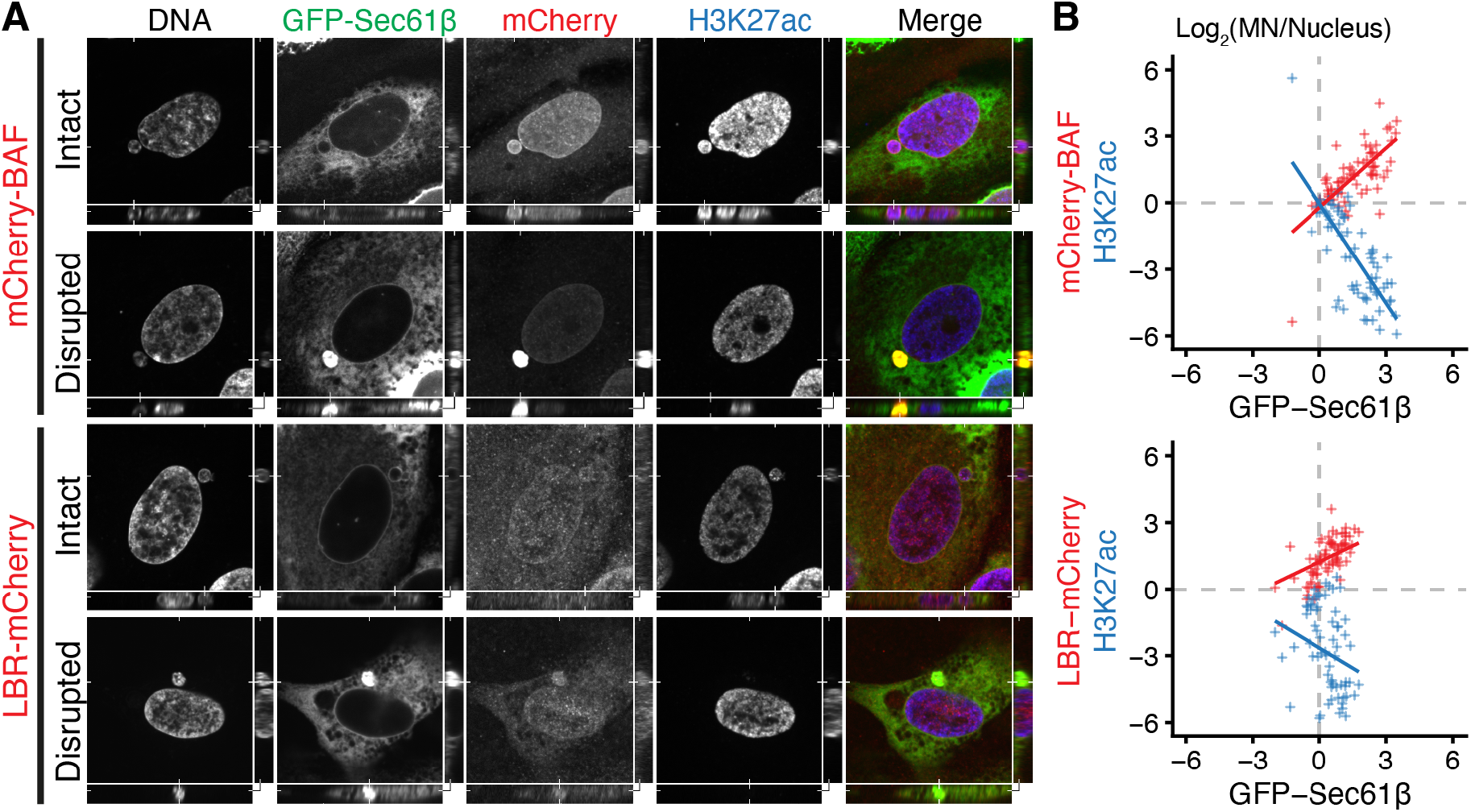
Missegregation of an ensheathed chromosome results in a micronucleus with a disrupted nuclear envelope. (**A**) Confocal images showing examples of intact or disrupted micronuclei as indicated. Images show mCherry-BAF or LBR-mCherry (red) stably co-expressed with GFP-Sec61β (green) in RPE-1 cells, H3K27ac was detected by immunofluorescence (blue), and DNA stained with DAPI. XY view is through the centre of the micronucleus, YZ (right) and XZ (below) are orthogonal views at the positions indicated. (**B**) Scatter plots to show the fluorescence intensity of H3K27ac (blue) and either mCherry-BAF or LBR-mCherry (red) versus GFP-Sec61β intensity. Data are plotted as the log2 ratio of intensity at the micronucleus versus main nucleus. For RPE1 GFP-Sec61β mCherry-BAF n=71 cells and LBR-mCherry n=73 cells, from 3 independent experiments in each cell type.

### Induced relocalization of ER enables the rescue of ensheathed chromosomes

Does ensheathing of misaligned chromosomes cause chromosome missegregation? To answer this question we sought a way to clear the mitotic ER and test if this enabled subsequent rescue of misaligned chromosomes to the metaphase plate. To clear the mitotic ER, we used an induced relocalization strategy (Figure 6A). Induced relocalization of small organelles has been demonstrated for Golgi, intracellular nanovesicles and endosomes, typically using heterodimerization of FKBP-rapamycin-FRB with the FKBP domain fused to the organelle and the FRB domain at the mitochondria (Dunlop et al., 2017; Hirst et al., 2015; Larocque et al., 2020; van Bergeijk et al., 2015). We reasoned that a large organellar network, such as the ER, may be cleared by inducing its relocalization to the cell boundary. Our strategy therefore comprised an ER-resident hook (FKBP-GFP-Sec61β) and a plasma membrane anchor (stargazin-mCherry-FRB) with application of rapamycin predicted to induce the relocalization of ER to the plasma membrane (Figure 6A). HCT116 cells were used for these experiments as they are near diploid, easy to transfect, and showed a similar fate and mitotic response to GSK923925 pre-treatment as RPE-1 (Supplementary Figure S5).

**Figure 6.**
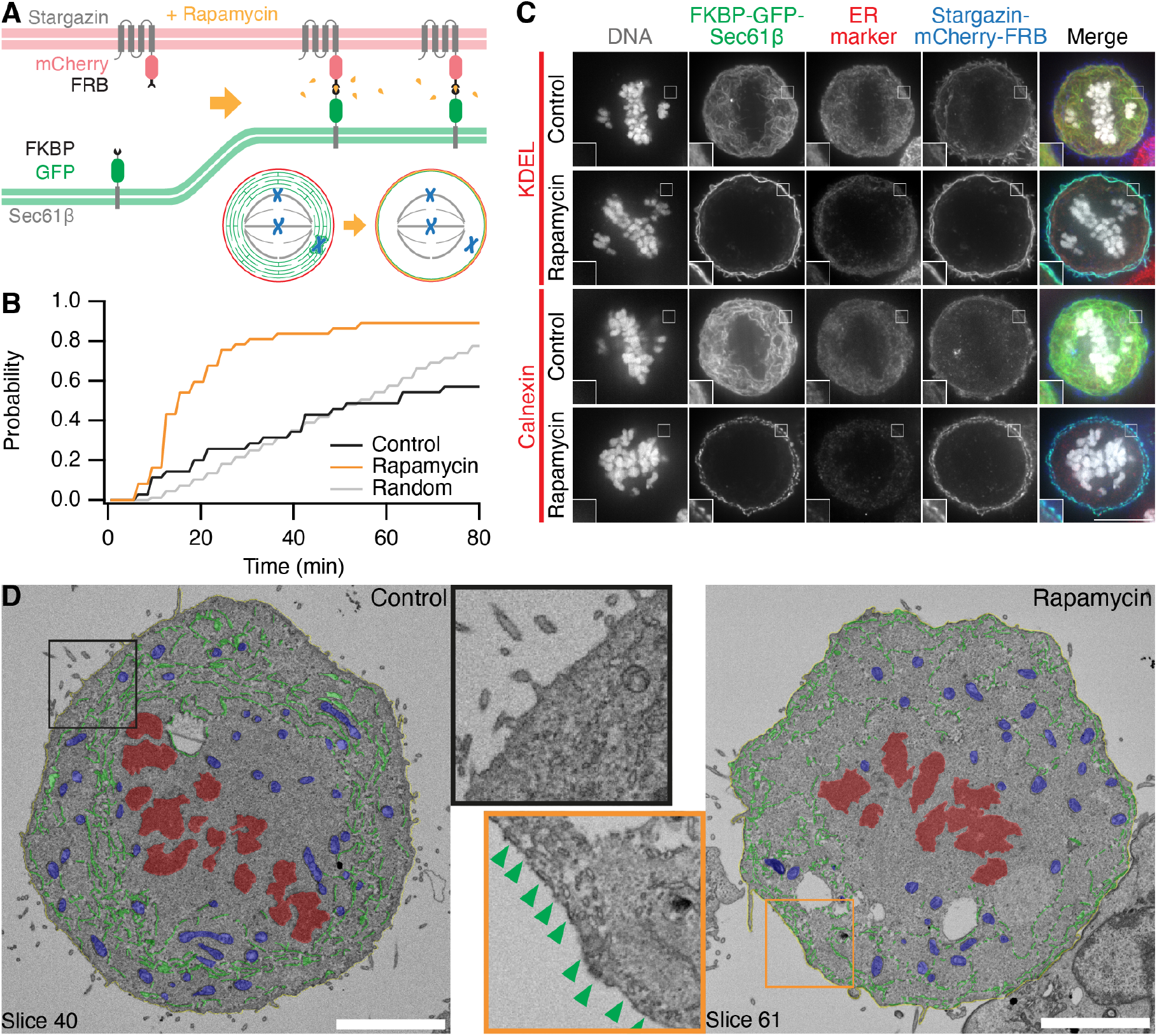
Inducible relocalization of ER in mitotic cells. (**A**) Schematic diagram of the ER clearance procedure. Rapamycin induces the heterodimerization of the ER-resident FKBP-GFP-Sec61β and the plasma-membrane localized Stargazin-mCherry-FRB. (**B**) Cumulative histogram showing the time to detection of ER clearance. An automated segmentation procedure was used to monitor ER localization in mitotic cells. The time at which the largest decrease in ER localization occurred was taken (see Methods). Random occurrence is shown for comparison. The median (IQR) ER clearance time in Rapamycin-treated cells was 15 (12–24) minutes, rapamycin is applied after the first frame (*T* = 0). (**C**) Induced relocalization of FKBP-GFP-Sec61β to the plasma membrane causes ER clearance. Typical immunofluorescence micrographs of mitotic HCT116 cells pre-treated with GSK923295, expressing FKBP-GFP-Sec61β (green) and Stargazin-mCherry-FRB (blue), treated or not with rapamycin (200 nM). Cells were stained for ER markers KDEL or Calnexin as indicated (red), DNA was stained with DAPI (gray). Insets are 2X expansions of the ROI shown. Scale bar, 10 µm. (**D**) Serial block face-scanning electron microscopy (SBF-SEM) imaging of control or ER-cleared (Rapamycin) mitotic HCT116 cells. A single slice is shown with segmentation of ER (green), plasma membrane (yellow), mitochondria (blue) and chromosomes (red). Scale bar, 5 µm. Insets are 2X expansions of the indicated ROI shown without segmentation; green arrowheads indicate ER attachment to the plasma membrane.

We found that the clearance of ER in mitotic cells with this strategy was efficient, occurring in 89.2 % of HCT116 cells expressing the system after treatment with 200 nM rapamycin. Onset was variable with a median time to maximum clearance of 15 min (IQR, 12 min to 24 min, Figure 6B). Importantly, induced relocalization of FKBP-GFP-Sec61β to the plasma membrane represented the clearance of ER and not the extraction of the protein. First, immunostaining of two other endogenous ER resident proteins KDEL and calnexin also showed relocalization to the plasma membrane (Figure 6C). Second, SBF-SEM imaging allowed us to observe the relocalization of ER to the plasma membrane (Figure 6D). Here, the expansion of the exclusion zone and the direct attachment of hundreds of ER tubules to the plasma membrane could be unambiguously visualized.

We next tested whether ER clearance could be used as an intervention in cells with ensheathed chromosomes. To do this, HCT116 cells expressing FKBP-GFP-Sec61β and stargazin-mCherry-FRB, pre-treated with 150 nM GSK923295 to induce ensheathed chromosomes, were imaged as 200 nM rapamycin was applied to induce clearance of the ER. In control cells where no rapamycin was applied, the cells were arrested in mitosis for prolonged periods. In cells where the ER had been cleared, congression of the ensheathed chromosome was clearly seen after clearance had occurred (Figure 7A and Supplementary Video SV7). We used automated image analysis to track the 3D position of the misaligned chromosome over time, in an unbiased manner (Figure 7B-C). Congression of the ensheathed chromosome within 80 min was seen in 86.7 % of cells with induced ER clearance. In control cells the majority (66.7 %) were unable to resolve the ensheathed chromosome in the same time (Figure 7A-C). These data suggest that ER clearance is an effective intervention in cells with ensheathed chromosomes and points to a causal role for endomembranes in chromosome missegregation.

**Figure 7.**
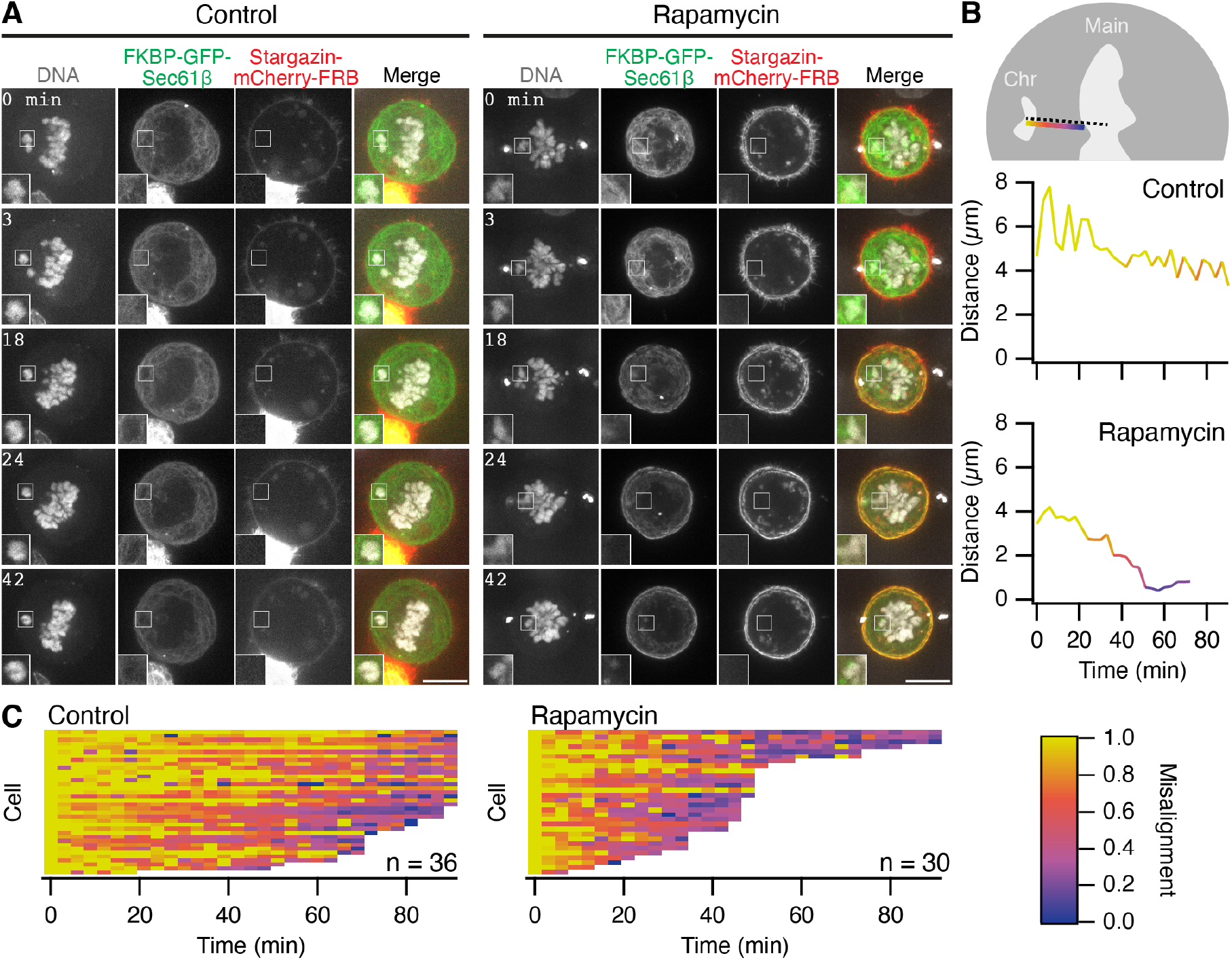
Rescue of ensheathed chromosomes by the induced relocalization of ER. (**A**) Stills from live cell imaging of ER clearance experiments. FKBP-GFP-Sec61β (green), Stargazin-mCherry-FRB (red) and SiR-DNA (gray) are shown. Insets are 2X expansions of the ROI shown. Scale bars, 10 µm. See Supplementary Videos SV7. (**B**) Semi-automated 4D tracking of misaligned chromosome location is used to monitor congression. Two tracks from the cells in A are shown. The shortest Euclidean distance from the centroid of the misaligned chromosome to the edge of the main chromosome plate is plotted as a function of time. (**C**) Fate of misaligned chromosomes in control or Rapamycin-treated cells. Rescue of misaligned chromosomes was detected in 26 out of 30 Rapamycin-treated cells. Coloring in B and C is with the colorscale shown. Tracks terminate at 90 min or when the chromosome merges with the plate. Median termination time was 93 min (Control) and 45 min (Rapamycin); p = 7.1 ×10^−9^, Wilcoxon Rank Test. Rapamycin is applied after the first frame (*T* = 0).

## Discussion

This study demonstrates that misaligned chromosomes located beyond the exclusion zone are liable to become ensheathed by endomembranes. The fate of cells with ensheathed chromosomes is biased towards missegregation, aneuploidy and micronuclei formation. We showed that if the ER was cleared by induced relocalization in live mitotic cells, these chromosomes could be rescued by the mitotic spindle, an intervention which suggests that chromosome ensheathing by endomembranes is a risk factor for chromosome missegregation and subsequent aneuploidy.

Chromosomes can become misaligned during mitosis for a number of reasons, but we show here that those that transit out of the exclusion zone become ensheathed in endomembranes. We demonstrated this with four different cell models: RPE-1 or HCT116 cells pre-treated with a CENP-E inhibitor, DLD-1 cells with targeted disconnection of the Y-chromosome and HeLa cells with spontaneously arising misaligned chromosomes. In each case misaligned chromosomes beyond the exclusion zone typically became ensheathed in endomembranes. Although the morphology of mitotic endomembranes varies between cell lines (Puhka et al., 2007; Lu et al., 2009, 2011; Puhka et al., 2012; Champion et al., 2017), all ensheathed chromosomes were draped in several layers of endomembranes. We use the term ensheathed to describe how these chromosomes are surrounded by endomembranes but not fully enclosed in any one layer as though in a vesicle. The ensheathing membrane follows the contours of the chromosome closely. Our SBF-SEM analysis did not uncover any obvious electron dense connections between the ensheathed chromosome and its surrounding membranes, although a previous report indicated that exogenous DNA clusters may physically interact with mitotic ER (Wang et al., 2016).

A major finding of our work is that ensheathing promotes missegregation and micronuclei formation. Our 3D EM images of ensheathed chromosomes show that microtubules face a difficult task to negotiate several layers of endomembranes in order to make the contact between kinetochore and spindle that is necessary for rescue and alignment. In cases where contact is made, endomembranes are also likely to impair the congression of the chromosome, as suggested by a recent study where excess ER was shown to slow chromosome motions (Merta et al., 2021). Since endomembranes are a risk factor for missegregation, their precise organization – for example the sheet-to-tubule ratio of the ER – may influence the likelihood for missegregation (Champion et al., 2017). The lack of attachment is sufficient to prolong spindle assembly checkpoint signalling and delay mitosis. Ultimately, the cells progress to anaphase and missegregate, likely due to checkpoint exhaustion after prolonged metaphase (Uetake and Sluder, 2010; Yang et al., 2008). Whatever the mechanism, the role of endomembranes in promoting missegregation may be important for tumor progression. It is possible that in tumor cells that are aneuploid, endomembranes may contribute to the higher rates of CIN observed (Funk et al., 2016; Nicholson and Cimini, 2015).

The fate of cells with ensheathed chromosomes was biased towards missegregation and formation of micronuclei. Interestingly, a previous study found that artificially tethering endomembranes to aligned chromosomes within the exclusion zone caused mitotic errors; although the outcome was dependent on at what stage tethering was induced (Champion et al., 2019). Tethering prior to mitotic entry resulted in segregation errors and multilobed nuclei, whereas tethering during metaphase had little consequence. Although conceptually similar, the ensheathing process reported here is a natural consequence of a misaligned chromosome becoming entangled in endomembranes. Key differences include the position of the ensheathed chromosome, the lack of microtubule attachments, no direct membrane-chromosome tethering, and multiple vs single endomembrane layers; these are likely to explain the different observed mitotic phenomena. We found that the micronuclei that result from ensheathed chromosomes had disrupted envelopes 8 h post-release from CENP-E inhibition. Rupture of micronuclei has been shown to lead to DNA damage, and activation of innate immune and cell invasion pathways (Ly et al., 2017; Hatch et al., 2013; Mam- mel et al., 2021; Bakhoum et al., 2018). The presence of ER in the micronuclear space of disrupted micronuclei indicates that ensheathing may increase the likelihood of rupture, we speculate that this may occur by endomembranes physically interfering with envelope reformation at the micronucleus.

Mitosis in human cells is open, yet we have known for over 60 years that the spindle exists in a membrane-free ellipsoid exclusion zone (Bajer, 1957; Porter and Machado, 1960; Nixon et al., 2017). It seems intuitive that the spindle must operate in a membrane-free area to avoid errors, but recent work suggests that the exclusion zone is actively maintained and that this arrangement is important for concentrating factors for spindle assembly (Schweizer et al., 2015) or for maintenance of spindle structure (Kumar et al., 2019; Schlaitz et al., 2013). We found that ER clearance, via an induced relocalization strategy, could be used as an intervention to improve the outcome for mitotic cells with ensheathed chromosomes. Induced relocalization of small organelles has previously been demonstrated (Dunlop et al., 2017; Hirst et al., 2015; Larocque et al., 2020; van Bergeijk et al., 2015), but the movement of a large organellar network by similar means had not been attempted previously. Surprisingly, ER clearance in mitotic cells was efficient, although it was much slower than the relocalization of intracellular nanovesicles, taking tens of minutes rather than tens of seconds (Larocque et al., 2020). We speculate that the efficiency of clearance is due to cooperativity of relocalization since the FKBP-GFP-Sec61β molecules are dispersed in the ER which is interconnected. These experiments were important to show that ensheathing was causal for chromosome missegregation. We note that this method has many future applications: to selectively perturb mitotic structures, at defined times, during cell division. For example, ER clearance and concomitant expansion of the exclusion zone is an ideal manipulation to probe the function of this enigmatic cellular region.

## Methods

### Molecular biology

The following plasmids were gifts, available from Addgene, or from previous work as indicated: Histone H3.2-mCherry (A. Bowman, University of Warwick), pAc-GFPC1-Sec61β (Addgene #15108), psPAX2 (Addgene #12260), pMD2.G (Addgene #12259), pWPT-GFP (Addgene #12255), Stargazin-GFP-LOVpep (Addgene #80406), LBR pEGFP-N1 (Addgene #61996), EGFP-BAF (addgene #101772), pMito-mCherry-FRB (Addgene #59352), Histone H2B-mCherry (Cheeseman et al., 2013), pFKBP-GFP-C1 (Clarke and Royle, 2018)

To generate a plasmid to express mCherry-Sec61β, EcoRI-BglII digestion product of pAc-GFPC1-Sec61β was ligated into pmCherry-C1 vector (made by substituting mCherry for EGFP in pEGFP-C1 (Clontech) by AgeI-XhoI digestion). LBR-mCherry was made by amplifying the LBR insert from LBR in pEGFP-N2 and ligating into pmCherry-N1 using KpnI and BamHI. The mCherry-BAF construct was amplified from EGFP-BAF and inserted into pmCherry-C1 using BglII and HindIII. For lentivirus transfer plasmids, constructs for expression (mCherry-BAF, GFP-Mad2, mCherry-Sec61β) were cloned into pWPT-GFP using MluI-SalI sites or MluI-BstBI for LBR-mCherry.

Plasmids for ER clearance were generated as follows. For FKBP-GFP-Sec61β, a BglII-EcoRI fragment from pAc-GFP-C1-Sec61β was ligated into pFKBP-GFP-C1. Stargazin-mCherry-FRB construct was made by PCR of Stargazin encoding region from Stargazin-GFP-LOVpep and insertion into pMito-mCherry-FRB at NheI-BamHI sites.

### Cell biology

HCT116 (ATCC, CCL-247) and HEK293T (ATCC, CRL-11268) cells were maintained in Dulbecco’s Modified Eagle’s Medium (DMEM) supplemented with 10 % FBS and 100 U ml^−1^ penicillin/streptomycin. DLD-1-WT and DLD-1-C-H3 (Ly et al., 2017) cell lines were gifts from Don Cleveland (UCSD). These cell lines and their derivatives were maintained in DMEM supplemented with 10 % Tetra-Free FBS (D2-118, SLS), 2 mM L-glutamine, 100 U ml^−1^ penicillin/streptomycin and 100 µg ml^−1^ hygromycin. RPE-1 (Horizon Discovery) and derived cell lines were maintained in DMEM/F-12 Ham supplemented with 10 % FBS, 2 mM L-glutamine, 100 U ml^−1^ penicillin/streptomycin and 0.26 % sodium bicarbonate (NaHCO_3_). All cell lines were kept in a humidified incubator at 37 ^*°*^C and 5 % CO_2_. Cells were routinely tested for mycoplasma contamination by a PCR-based method.

RPE-1 GFP-Sec61β stable cell line was generated by Fugene-HD (Promega) transfection of pAc-GFPC1-Sec61β. DLD-1-WT mCherry-Sec61β and DLD-1-C-H3 mCherry-Sec61β stable cell lines were generated by GeneJuice (Merck Millipore) transfection of mCherry-Sec61β into the respective parental lines. Individual clones were isolated by G418 treatment (500 µg ml^−1^) and were validated using a combination of western blot, FACS and fluorescence microscopy. Stable co-expression of Histone H3.2-mCherry, mCherry-BAF or LBR-mCherry with GFP-Sec61β in RPE-1 cells was achieved by lentiviral transduction of cells stably expressing GFP-Sec61β. For stable expression of GFP-Mad2 with mCherry-Sec61β, dual lentivirus transduction was used. Individual cells positive for GFP and mCherry signal were sorted by FACS and single cell clones validated by fluorescence microscopy. Note that the transgenic expression of GFP-Sec61β is associated with downregulation of endogenous Sec61β (Supplementary Figure S3A). Transient transfections of HCT116, RPE-1 and HeLa were done using Fugene-HD or GeneJuice according to the manufacturer’s instructions.

For lentiviral transduction, HEK293T packaging cells were incubated in DMEM supplemented with 10 % FBS, 2 mM L-glutamine, and 25 µM chloroquine diphosphate (C6628, Sigma) for 3 h. Transfection constructs were prepared at 1.3 pM psPAX2, 0.72 pM pMD2.G and 1.64 pM transfer plasmid (encoding the tagged protein to be expressed) in OptiPro SFM. Polyethylenimine (PEI) dilution in OptiPro SFM was prepared separately at 1:3 ratio with DNA (w/w, DNA:PEI) in the transfection mixture. Transfection mixes were combined, incubated at room temperature for 15 min to 20 min, and then added to the packaging cells. Cells were incubated for 18 h, after which the media was replaced with DMEM supplemented with 10 % FBS and 100 U ml^−1^ penicillin/streptomycin. Viral particles were harvested 48 h post-transfection. Viral supernatant was centrifuged and filtered before applying to target cells. Target cells were then infected through incubation in media containing 8 µg ml^−1^ polybrene (408727, Sigma) for 16 h to 20 h. Media was replaced with complete media and cells were screened after 24 h. All incubations were in a humidified incubator at 37 ^*°*^C and 5 % CO_2_.

To induce misaligned chromosomes in RPE-1 or HCT116 cell lines, cells were incubated in complete media containing 150 nM GSK923295 (Selleckchem) for 3 h before release of cells from treatment. For fixed cell experiments, release was for 1 h. To induce the auxin-degron system in DLD-1 cells, 500 µM indole-3-acetic (A10556, Fisher) and 500 µg ml^−1^ doxycycline (Sigma, D9891) were added to the media and cells incubated for 24 h.

ER clearance was induced through application of rapamycin (Alfa Aesar) to final concentration of 200 nM, to HCT116 cells expressing FKBP-GFP-Sec61β and stargazin-mCherry-FRB. For fixed cell experiments, rapamycin treatment was for 30 min.

### Fluorescence methods

For immunofluorescence, cells were fixed at room temperature using PFA solution (3 % formaldehyde, 4 % sucrose in phosphate-buffered saline (PBS)) for 15 min and permeabilized at room temperature in 0.5 % (v/v) Triton X-100 in PBS for 10 min. Cells were blocked in 3 % BSA in PBS for 60 min at room temperature. Cells were then incubated for 60 min at room temperature with primary antibody dilutions prepared in 3 % BSA in PBS as follows: mouse anti-Bub1 (ab54893, Abcam, 1:500); mouse anti-Mad2 (sc-65492, Santa Cruz, 1/200); rabbit anti-calnexin (ab22595, Abcam, 1:200); guinea pig anti-CENP-C (PD030, Medical and Biological Labs Company, 1:2000); rabbit anti-H3K27ac (ab4729, Abcam, 1:1000); rabbit anti-KDEL (PA1-013, Invitrogen, 1:200); rabbit anti-kinastrin (HPA042027, Atlas Antibodies, 1:1000). Following three PBS washes, cells were incubated with secondary antibodies for 60 min, either Alexa Fluor568- or Alexa Fluor647-conjugated antibody in 3 % BSA/PBS (Invitrogen, 1:500). Following three PBS washes, coverslips were rinsed and mounted with Vectashield containing 4’,6-diamidino-2-phenylindole (DAPI, Vector Laboratories), then sealed. In cases where GFP signal required amplification, cells were incubated with GFP-booster (Alexa Fluor488, Chromotek, 1:200) at the secondary antibody step. Where amplification of mCherry was required, mouse anti-mCherry (1C51 - ab125096, Abcam, 1:500) was used with AlexaFluor568-conjugated secondary antibody.

For fluorescent *in situ* hybridization (FISH) of DLD-1 WT and DLD-1-C-H3 cells, the degron system was induced and cells synchronized by doubled thymidine (2.5 mM) treatment. Samples were fixed in Carnoy’s fixative (3:1 v/v methanol:glacial acetic acid) for 5 min at room temperature, then rinsed in fixative, before addition of fresh fixative and incubated for a further 10 min. Samples were rinsed in distilled water before FISH probe denaturation and hybridization following the manufacturer’s protocol (Xcyting Centromere Enumeration Probe, XCE Y green, D-0824-050-FI, MetaSystems Probes).

To dye chromosomes or microtubules in fixed or live cell imaging, cells were incubated for 30 min with 0.5 µM SiR-DNA or SiR-Tubulin (Spirochrome), respectively.

### Biochemistry

For western blot, cells were harvested, and lysates prepared by sonication of cells in UTB buffer (8 M Urea, 50 mM TRIS, 150 mM β-mercaptoethanol). Lysates were incubated on ice for 30 min, clarified in a benchtop centrifuge (20 800 *g*) for 15 min at 4 ^*°*^C, then boiled in Laemmli buffer for 10 min and resolved on a precast 4 % - 15 % polyacrylamide gel (Bio-Rad). Proteins were transferred to nitrocellulose using a Trans-Blot Turbo Transfer System (Bio-Rad). Primary antibodies were diluted in 4 % BSA in PBS and used as follows: rabbit anti-Sec61β (PA3-015, Invitrogen, 1:1000); HRP conjugated mouse anti-β-actin (sc-47778, Santa Cruz, 1: 20000); rabbit anti-mCherry (ab183628, Abcam, 1:2000); anti-GAPDH (G9545, Sigma, 1/5000); rabbit anti-CENP-A (2186, Cell Signalling, 1:1000); mouse anti-BAF (A-11, Santa Cruz, 1:500); mouse anti-LBR (SAB1400151, Sigma-Aldrich, 1:500). Secondary antibodies of anti-mouse, -rabbit and -rat IgG HRP conjugates were prepared in 5 % milk in PBS. For detection, enhanced chemiluminescence detection reagent (GE Healthcare) and manual exposure of Hyperfilm (GE Healthcare) was performed.

### Microscopy

For fixed cell imaging experiments, a Personal DeltaVision microscope system (Applied Precision, LLC), based on an IX-71 microscope body (Olympus) was used with a CoolSNAP HQ2 interline CCD camera (Photometrics) and a 60× oil-immersion 1.42 NA oil PlanApo N objective. Equipped with Precision Control microscope incubator, Tokai Hit stage top incubator and Applied Precision motorized xyz stage. Illumination was via a Lumencor SPECTRA X light engine (DAPI, 395/25; GFP, 470/24; mCherry, 575/25; CY-5, 640/30), dichroics (Quad: Reflection 381-401:464-492:531-556:619-644 Transmission 409-456:500-523:564-611:652-700; GFP/mCh: Reflection 464-492:561-590 Transmission 500-553:598-617) and filter sets (DAPI: ex 387/11 em 457/50; GFP: ex 470/40 em 525/50; TRITC: ex 575/25 em 597/45; mCherry: ex 572/28 em 632/60; CY-5: ex 640/14 em 685/40). Image capture was by softWoRx 5.5.1 (Applied Precision). Images were deconvolved using softWoRx 3.0 with the following settings: conservative ratio, 15 cycles and high noise-filtering.

For live cell imaging, cells were plated onto fluorodishes (WPI) and imaged in complete media in an incubated chamber at 37 ^*°*^C and 5 % CO_2_. Most live cell imaging was done using a Nikon CSU-W1 spinning disc confocal system with SoRa upgrade (Yokogawa) was used with either a Nikon, 100x, 1.49 NA, oil, CFI SR HP Apo TIRF or a 63×, 1.40 NA, oil, CFI Plan Apo objective (Nikon) with optional 2.3× intermediate magnification and 95B Prime camera (Photometrics). The system has CSU-W1 (Yokogawa) spinning disk unit with 50 µm and SoRa disks (SoRa disk used), Nikon Perfect Focus autofocus, Okolab microscope incubator, Nikon motorized xy stage and Nikon 200 µm z-piezo. Excitation was via 405 nm, 488 nm, 561 nm and 638 nm lasers with 405/488/561/640 nm dichroic and Blue, 446/60; Green, 525/50; Red, 600/52; FRed, 708/75 emission filters. Acquisition and image capture was via NiS Elements (Nikon).

For mitotic progression and fate experiments, the DeltaVision system described above was used. For live cell imaging of HeLa cells, a spinning disc confocal system (Ultra-View VoX, PerkinElmer) with a 60x, 1.40 NA, oil, Plan Apo VC objective (Nikon) was used. Images were captured using an ORCA-R2 digital charge-coupled device camera (Hamamatsu) following excitation with 488 nm and 561 nm lasers, and 405/488/561/640 nm dichroic and 525/50, 615/70 filter sets. Images were captured using Volocity 6.3.1.

All microscopy data was stored in an OMERO database in native file formats.

### Serial block face-scanning electron microscopy

To prepare samples for serial block face-scanning electron microscopy (SBF-SEM), RPE-1 GFP-Sec61β on gridded dishes were first incubated with 150 nM GSK923295 (Selleckchem) for 3 h to induce misaligned chromosomes, before release of cells from treatment and incubation for around 30 min with 0.5 µM SiR-DNA (Spirochrome) to visualize DNA. HeLa cells on gridded dishes were not treated and were not stained. Using live cell light microscopy, cells with an ensheathed chromosome were selected for SBF-SEM. Fluorescent and bright-field images of the selected cell were captured and the coordinate position recorded. Cells were washed twice with PB (phosphate buffer) before fixing (2.5 % glutaraldehyde, 2 % paraformaldehyde, 0.1 % tannic acid (low molecular weight) in 0.1 M phosphate buffer, pH 7.4) for 1 h at room temperature. Samples were washed three times with PB and then post-fixed in 2 % reduced osmium (equal volume of 4 % OsO_4_ prepared in water and 3 % potassium ferrocyanide in 0.1 M PB solution) for 1 h at room temperature, followed by a further three washes with PB. Cells were then incubated for 5 min at room temperature in 1 % (w/v) thiocarbohydrazide solution, followed by three PB washes. A second osmium staining step was then included, incubating cells in a 2 % OsO_4_ solution prepared in water for 30 min at room temperature, followed by three washes with PB. Cells were then incubated in 1 % uranyl acetate solution at 4 ^*°*^C overnight. This was followed by a further three washes with PB. Walton’s lead aspartate was prepared adding 66 mg lead nitrate (TAAB) to 9 ml 0.03 M aspartic acid solution at pH 4.5, and then adjusting to final volume of 10 ml with 0.03 M aspartic acid solution and to pH 5.5 (pH adjustments with KOH). Cells were incubated in Walton’s lead aspartate for 30 min at room temperature and then washed three times in PB. Samples were dehydrated in an ethanol dilution series (30 %, 50 %, 70 %, 90 %, and 100 % ethanol, 5 min incubation in each solution) on ice, then incubated for a further 10 min in 100 % ethanol at room temperature. Finally, samples were embedded in an agar resin (AGAR 100 R1140, Agar Scientific).

SBF-SEM data was segmented using Microscopy Image Browser version 2.60 and the resulting 3D model was visualized in IMOD version 4.10.49 (Belevich et al., 2016; Kremer et al., 1996). HeLa SBF-SEM data was segmented and reconstructed in Amira 6.7 (Thermo Fisher Scientific).

### Data analysis

Kinetochore position analysis was in two parts. First, the position of kinetochores and spindle poles in hyperstacks were manually mapped using Cell Counter in Fiji. The kinetochore pointsets were classified into three categories: those aligned at the metaphase plate and those that were misaligned, with this latter group subdivided into kinetochores of chromosomes that were ensheathed and those that were not (free). Second, the ER channel of the hyperstack was segmented in Fiji to delineate the exclusion zone. Next, the Cell Counter XML files and their respective binarized ER stacks were read by program written in Igor Pro (WaveMetrics). To analyze the position of points relative to the exclusion zone in each cell, the ratio of two Euclidean distances was calculated (see Equation 1). Where *C* is the centroid of all aligned kinetochores, *Pi* is the position of a kinetochore and *Qi* is the point on the path from *C* through *P*, where the exclusion zone/ER boundary intersects with the path.

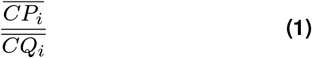

The ratio of these two distances gave a measure of how deep the point was placed inside or outside the exclusion zone (0 being on the boundary and 1 being as far outside of the exclusion zone as from the centroid to the boundary, on a log2 scale).

For analysis of live cell GFP-Mad2 and mCherry-Sec61β imaging, a semi-automated 4D tracking procedure was used. Briefly, the DNA channel from these movies was used for segmentation of chromosomes and metaphase plate as discrete 3D objects over time. The centroid-to-centroid distance was found for each chromosome relative to the plate (congression was taken as the merging of chromosome and plate objects) and the time of anaphase onset determined. Fluorescence signals were taken from each chromosome object using a 3 pixel expansion of the ROI. For mCherry-Sec61β, the mean voxel density was used. For GFP-Mad2, the maximum pixel intensity at each z position was taken from the expanded ROI and averaged per time point; this method gave a more accurate measure of Mad2 recruitment than the mean voxel density. Signals from each channel are expressed as a ratio of chromosome to plate. Mad2 signals were grouped by whether the chromosome congressed or not and then measurements from all chromosomes relative to anaphase were used to fit a line by linear regression. Only the last chromosome to congress (or not) was analyzed per cell. Data processing was via Fiji/ImageJ followed by analysis in Igor Pro.

Automated kinetochore-kinastrin colocalization was using a script that located the 3D position of kinetochores (CENP-C) and kinastrin puncta from thresholded images using 3D Object Counter in Fiji. These positions were loaded into Igor and the Euclidean distance to the nearest kinastrin punctum from each kinetochore was found.

ER clearance experiments were quantified using two automated procedures. First, ER, DNA and plasma membrane were segmented separately and then the plasma membrane segments were used to define the cell and the total area of segmented ER within this region was measured for all z-positions over time using a FIJI macro. Data were read by Igor and the ER volume over time was calculated. ER clearance manifested as a rapid decrease in ER volume, but the onset was variable. The derivative of ER volume of time was used to find the point of rapid decrease and this point was used to define the time to ER clearance. Random fluctuations in otherwise constant ER volume over time also resulted in a minima that occurred randomly. This process was modeled and plotted for comparison with the control group, where no clearance was seen. Second, the segmented DNA was classified into misaligned chromosome and main chromosome mass by a user blind to the conditions of the experiment. 3D coordinates of these two groups were fed into Igor where the centroids and boundaries of the chromosome and main chromosome mass where defined. The closest Euclidean distance between the centroid of the chromosome and edge of the main chromosome mass was used as the distance. Misalignment, shown as a colorscale, is this distance normalized to the starting distance.

Figures were made with FIJI, R or Igor Pro, and assembled using Adobe Illustrator. Null hypothesis statistical tests were done as described in the figure legends.

## Data and software availability

All code used in the manuscript is available at https://github.com/quantixed/Misseg

## ACKNOWLEDGEMENTS

We thank Claire Mitchell and Laura Cooper from the Computing and Advanced Microscopy Unit (CAMDU) for their help and support. Faye Nixon, Alison Beckett and Ian Prior at the Liverpool Biomedical EM Unit provided SBF-SEM imaging. We are grateful to Richard Bayliss, Vishakha Karnawat, Jonathan Millar, Alonso Pardal and James Shelford for feedback on the project and manuscript. This work was supported by a Pioneer Award and Programme Award from Cancer Research UK (C25425/A24167 and C25425/A27718). LD was supported by a studentship from BBSRC Midlands Integrative Biosciences Training Partnership (MIBTP) (BB/M01116X/1).

## AUTHOR CONTRIBUTIONS

NF, conceptualization, investigation, resources, supervision, writing - original draft, writing - review & editing. LD, formal analysis, investigation, resources, writing - review & editing. GS, visualization. SJR, formal analysis, funding acquisition, software, supervision, visualization, writing - original draft, writing - review & editing.

## COMPETING FINANCIAL INTERESTS

The authors declare no conflict of interest.

## Supplementary Information

**Figure S1.**
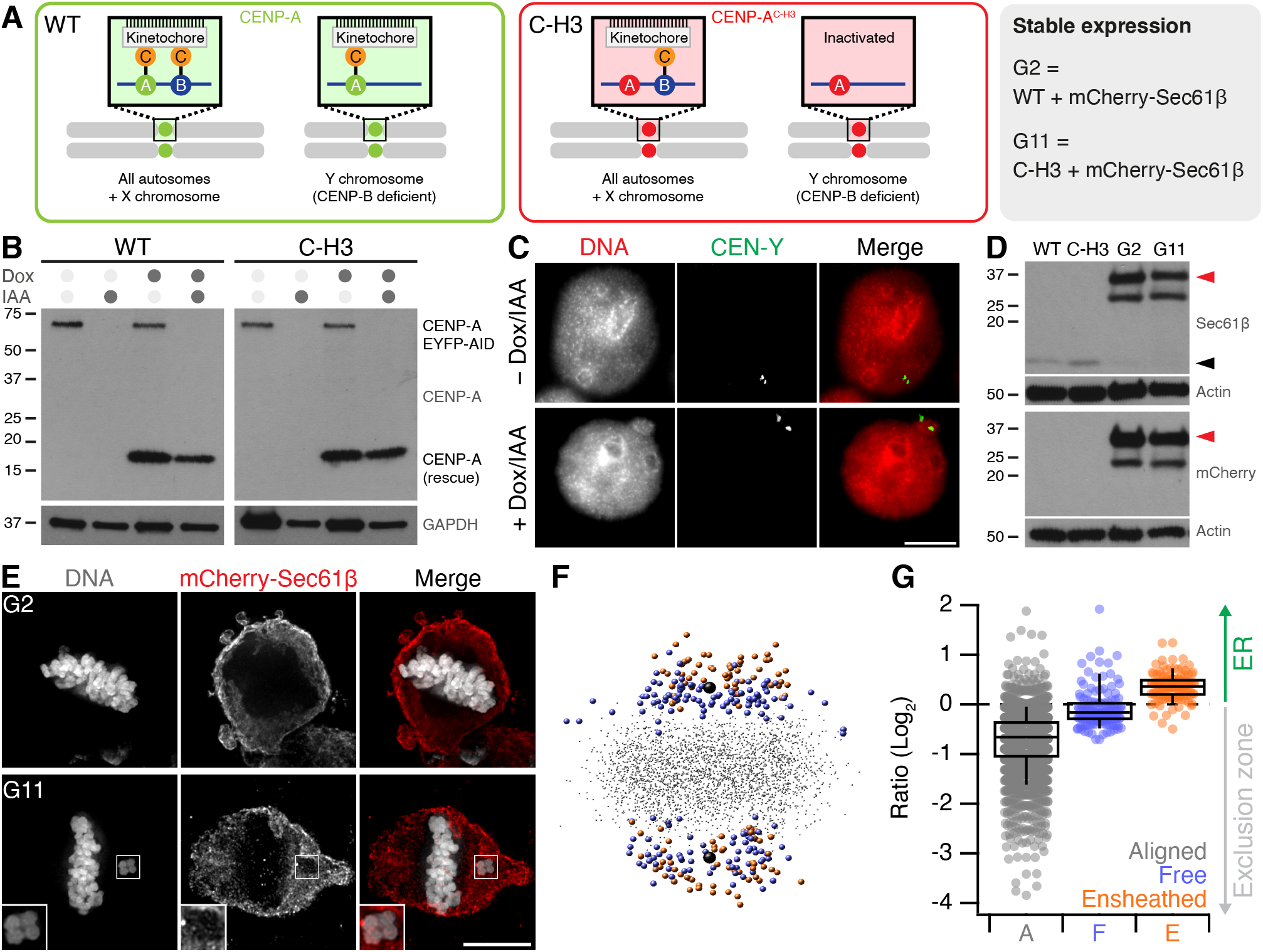
Ensheathed chromosomes in DLD-1 cells after targeted missegregation of Y-chromosome. (**A**) Schematic diagram after Ly et al. 2017, showing how re-expression of a CENP-A mutant (C-H3) in DLD-1 cells where CENP-A is degraded causes selective misalignment of the Y-chromosome. WT and C-H3 lines were further modified to express mCherry-Sec61β. (**B**) Western blot of lysates from WT or C-H3 DLD-1 cells treated with doxycycline (Dox) and/or indole-3-acetic acid (IAA) as indicated. Upper blot shows anti-CENP-A detection of endogenous CENP-A fused to EYFP-AID tag (66 kDa and expression of untagged CENP-A (either WT or C-H3). Lower blot shows GAPDH loading control. (**C**) Typical FISH images locating the Y-chromosome in the main nucleus in control cells and in a micronucleus in cells expressing C-H3 CENP-A. (**D**) Western blot of lysates from stable cell lines expressing mCherry-Sec61β derived from WT (G2) or C-H3 (G11). Detection of Sec61β or mCherry is shown as indicated with actin loading controls. Migration of Sec61β and mCherry-Sec61β are indicated by black and red arrowheads respectively. Note the expression of mCherry-Sec61β downregulates endogenous Sec61β. (**E**) Deconvolved widefield microscopy images showing an ensheathed chromosome in G11 cells but not in G2 cells treated with Dox/IAA. (**F**) Spatially averaged view of all kinetochores in the G11 DLD-1 Dox/IAA dataset (see Methods). Small gray points represent kinetochores at the metaphase plate. Colored points represent misaligned chromosomes that were ensheathed (orange) and those that were not (blue). Spindle poles are shown in black. (**G**) Box plot to show the relative position of each kinetochore relative to the exclusion zone boundary. Ratio of kinetochores within the exclusion zone are *<* 0 and those within the ER are *>* 0 on a log2 scale. Dots represent kinetochore ratios from 50 of DLD-1 cells at metaphase. Boxes show IQR, bar represents the median and whiskers show 9th and 91st percentiles.

**Figure S2.**
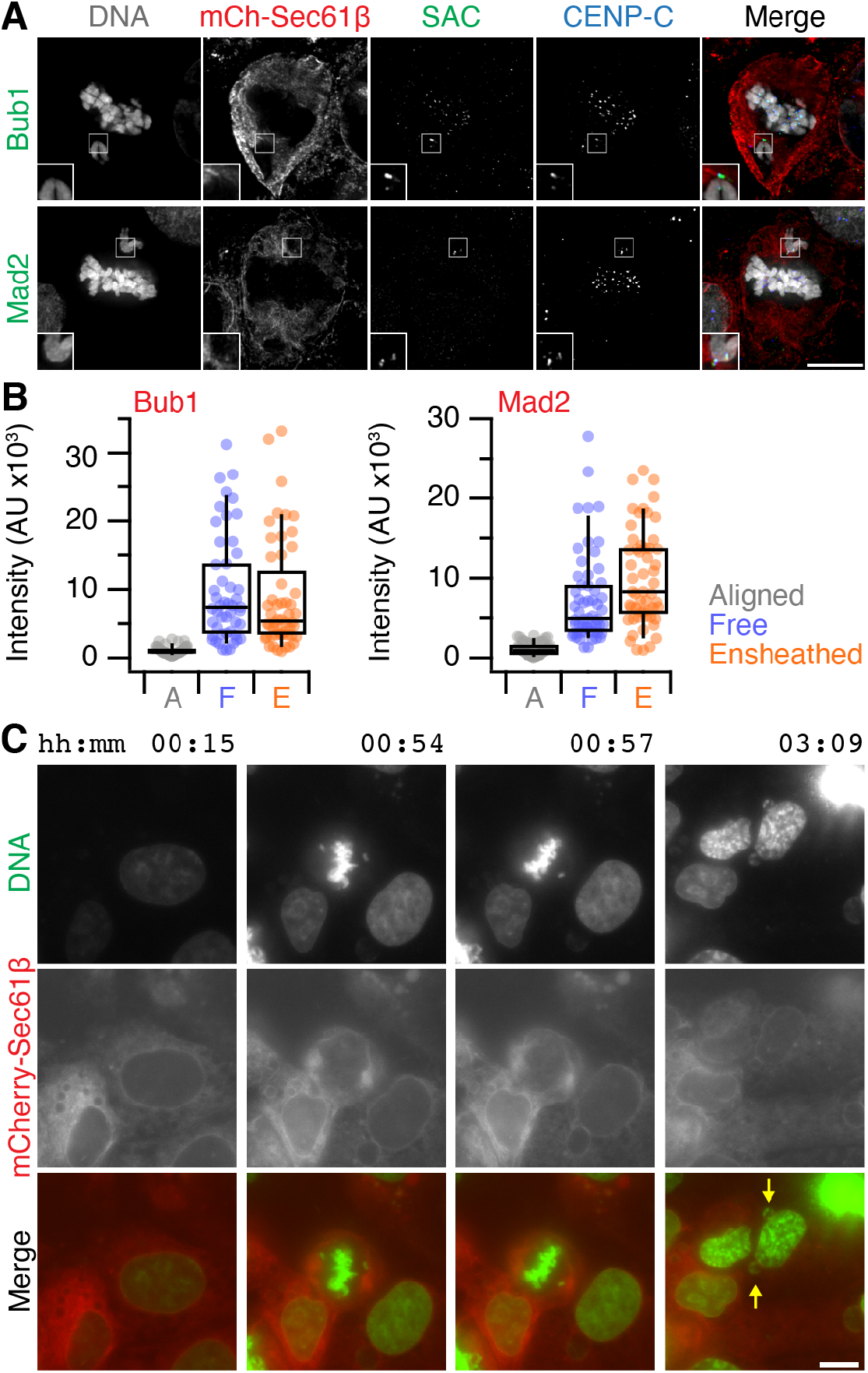
Spindle assembly checkpoint and micronuclei formation in DLD-1 cells. (**A**) Micrographs of immunofluorescence experiments to detect Bub1 or Mad2 (SAC, Green) at kinetochores (CENP-C, blue) in cells stably expressing mCherry-Sec61β (red), DAPI-stained DNA is shown in gray. Scale bar, 10 µm. (**B**) Quantification of Bub1 and Mad2 immunofluorescence at kinetochores marked by CENP-C. Ensheathed chromosomes were classified using the mCherry-Sec61β signal. (**C**) Stills from a movie showing an example of ensheathed chromosomes in G11 DLD-1 cells forming micronuclei following Dox/IAA treatment. Scale, 10 µm.

**Figure S3.**
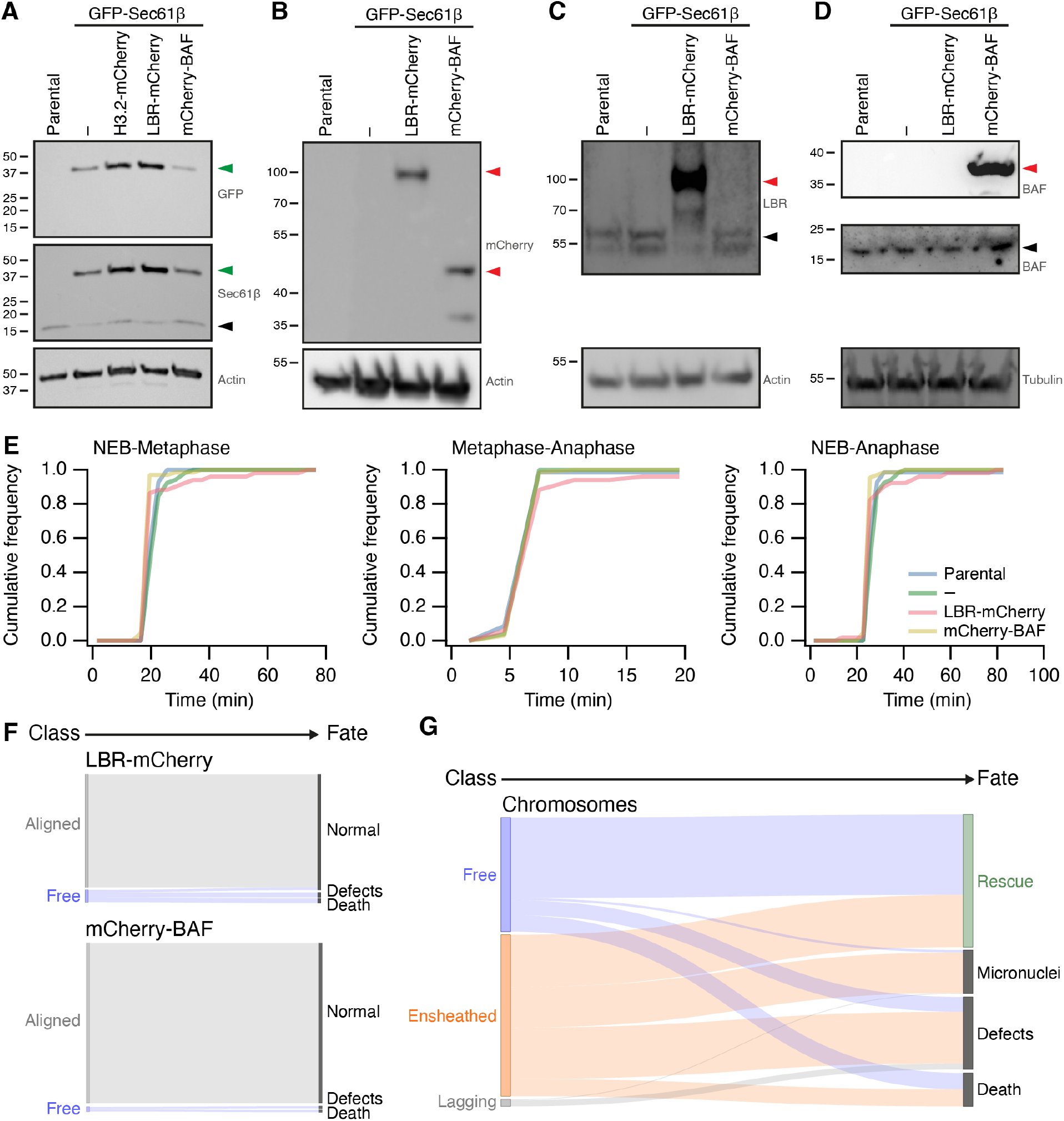
Stable transgene expression in RPE1 cells and fate of misaligned chromosomes in RPE1 cells stably expressing GFP-Sec61β. (**A-D**) Western blots to examine expression of proteins in parental RPE1 cells or clonal cells stably expressing GFP-Sec61β alone or with either Histone3.2-mCherry, LBR-mCherry or mCherry-BAF, as indicated. Membranes were probed for GFP, Sec61β, mCherry, LBR, BAF. Actin or tubulin is shown as a loading control. Green or red arrowheads indicate the expected position of GFP- or mCherry-tagged protein, black arrowheads indicate the untagged protein. (**E**) Mitotic timing of RPE1 cells stably expressing transgenes. Cumulative frequencies for nuclear envelope breakdown to metaphase, metaphase to anaphase and NEB to anaphase are shown. Parental, *n* = 69, GFP-Sec61β alone, *n* = 52, GFP-Sec61β and LBR-mCherry, *n* = 66, GFP-Sec61β and mCherry-BAF, *n* = 51. (**F**) Sankey diagram to show the fate (right) of RPE1 cells in each of the three metaphase classes (left). Fates include normal division, micronuclei formation, death and other defects (lagging chromosome, cytokinesis failure). Note that the fate of cells (and not chromosomes) is tracked. LBR-mCherry/GFP-Sec61β, *n* = 51, mCherry-BAF/GFP-Sec61β, *n* = 67; pooled from three experiments. (**G**) Sankey diagram to show the fate (right) of *chromosomes* in each of the three metaphase classes (left) after GSK923295 pre-treatment. Fates include rescue, micronuclei formation, death and other defects (lagging chromosome, cytokinesis failure). Number of chromosomes: Free, 146; Ensheathed, 207; Lagging, 9. The same dataset was analyzed for the outcome of *cells* (classified by the final misaligned chromosome) in Figure 4. Note that ensheathed chromosomes at metaphase that were rescued all became “free” chromosomes prior to rescue.

**Figure S4.**
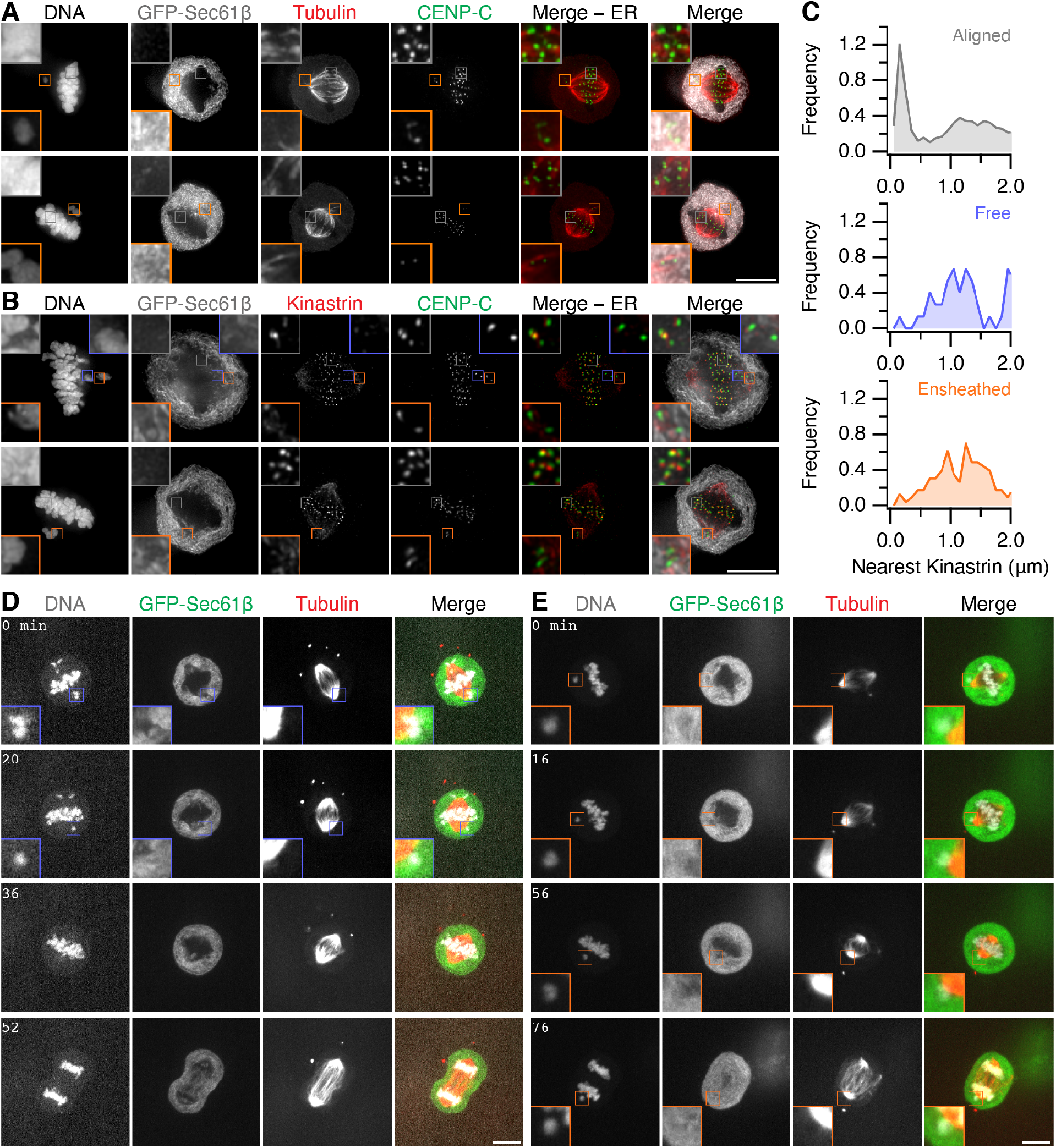
Ensheathed chromosomes do not have stable microtubule-kinetochore attachment. (**A**) Micrographs of RPE-1 cells stably expressing GFP-Sec61β (gray) pre-treated with GSK923295 immunostained for tubulin (red) and CENP-C (green), DNA stained with DAPI. Examples show end-on attachments at aligned kinetochores, and potential lateral kinetochore-MT contacts for ensheathed chromosomes. (**B**) Micrographs of RPE-1 cells stably expressing GFP-Sec61β (gray) pre-treated with GSK923295 immunostained for kinastrin (red) and CENP-C (green), DNA stained by DAPI. Scale bar, 10 µm. (**C**) Frequency distributions of the proximity of the nearest kinastrin punctum to each kinetochore (CENP-C punctum). Kinetochores (n, % with kinastrin <600 nm): aligned (3124, 26.8 %); free (74, 4.1 %); ensheathed (227, 6.2 %). (**D-E**) Still images from live-cell imaging experiments of RPE-1 cells stably expressing GFP-Sec61β (green) and Histone H3.2-mCherry (gray), pretreated with 150 nM GSK923295 and stained with SiR-Tubulin (Tubulin, red). Similar results were recorded in 25 cells with free chromosomes and 16 cells with ensheathed chromosomes. Inset shows a 2X zoom of the indicated region. Scale bar, 10 µm.

**Figure S5.**
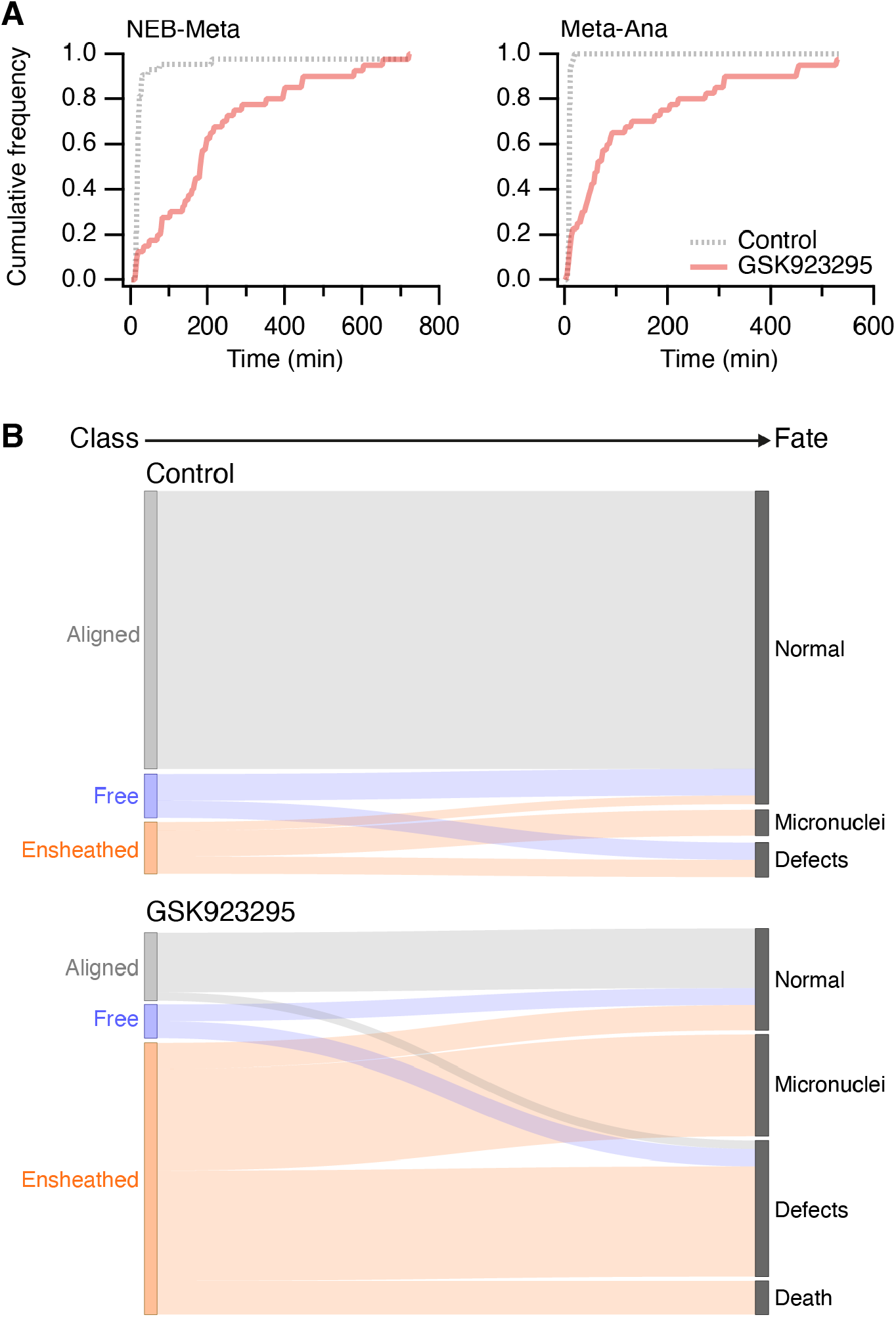
Mitotic timing and fate of HCT116 cells pre-treated with CENP-E inhibitor. (**A**) Mitotic timing of HCT116 cells. Cumulative frequencies for nuclear envelope breakdown to metaphase (NEB-Meta) and metaphase to anaphase (Meta-Ana) are shown. Cells were treated with 150 nM GSK923295 for 3 h before washout for 1 h and subsequent imaging. Control, *n* = 43, GSK pre-treatment, *n* = 40; pooled from three experiments. (**B**) Sankey diagram to show the fate (right) of cells in each of the three metaphase classes (left). Fates include normal division, micronuclei formation, death and other defects (lagging chromosome, cytokinesis failure). Note that the fate of cells (and not chromosomes) is tracked.

## Supplementary Videos

**Figure SV1.**
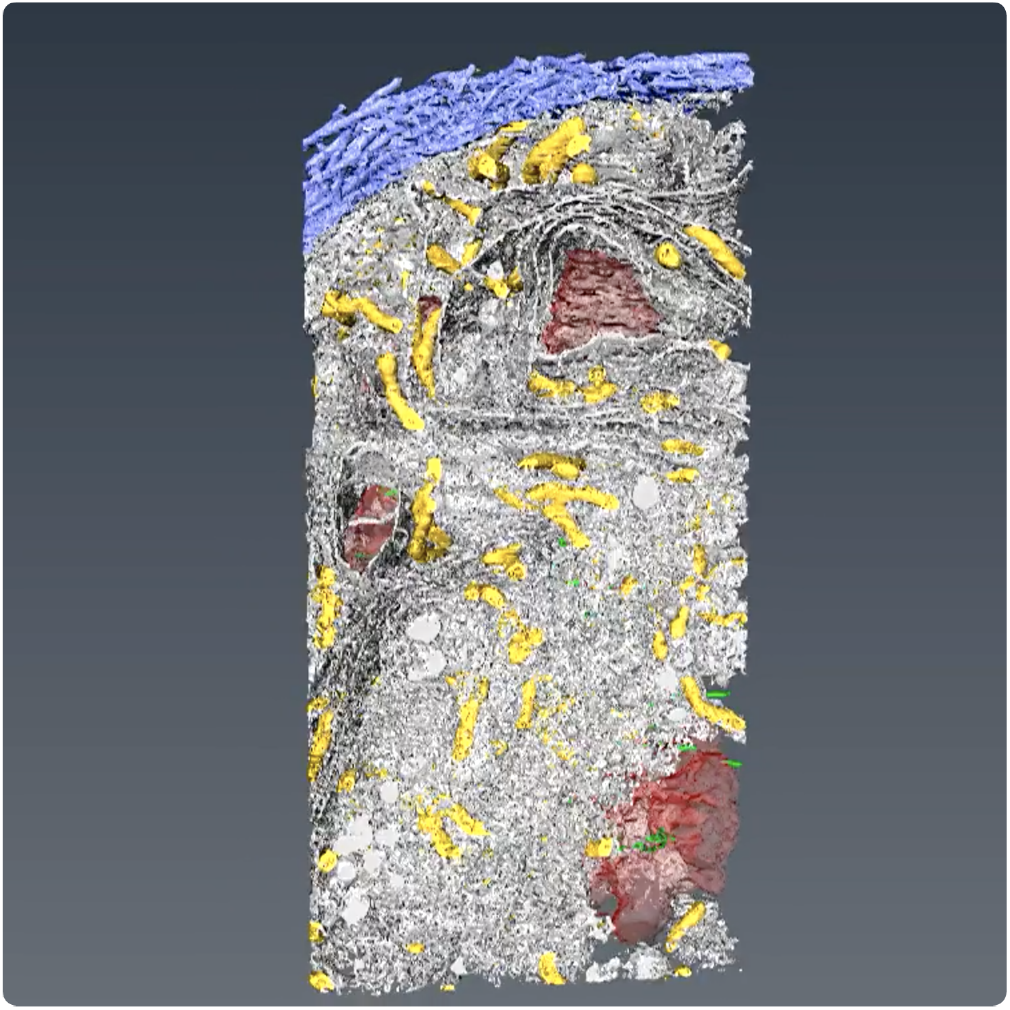
3D reconstruction of an ensheathed chromosome in a HeLa cell. SBF-SEM data from a HeLa cell with spontaneously occurring ensheathed chromosome. The following cellular features are shown (in order of appearance: spindle microtubules (green), centrioles (yellow), DNA (red), mitochondria (multi-colored then gold), endomembranes (white), plasma membrane (blue).

**Figure SV2.**
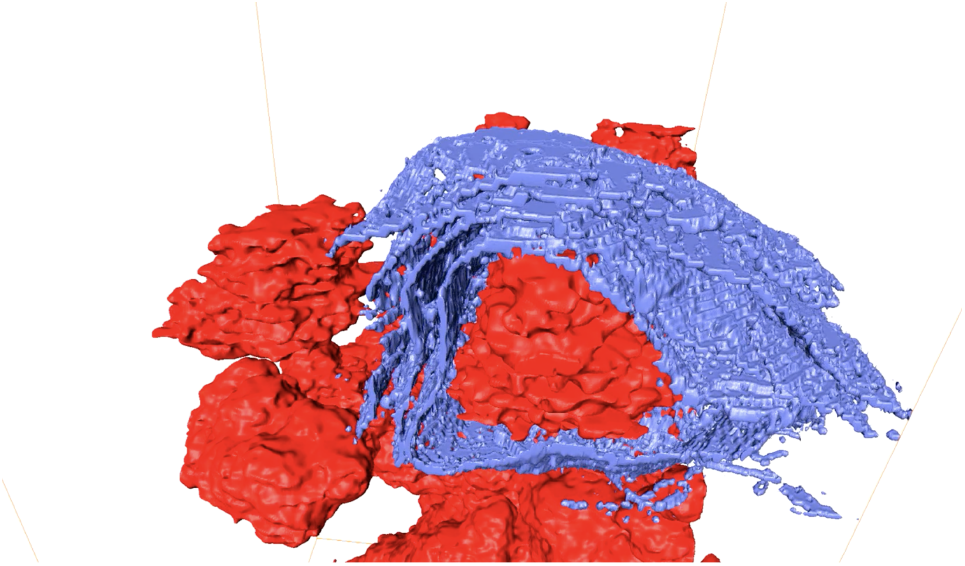
3D reconstruction of an ensheathed chromosome in a HeLa cell. Same reconstruction but showing only chromosomes (red) and endomembranes (blue). Endomembranes that ensheath the chromosome of interest are shown in purple.

**Figure SV3.**
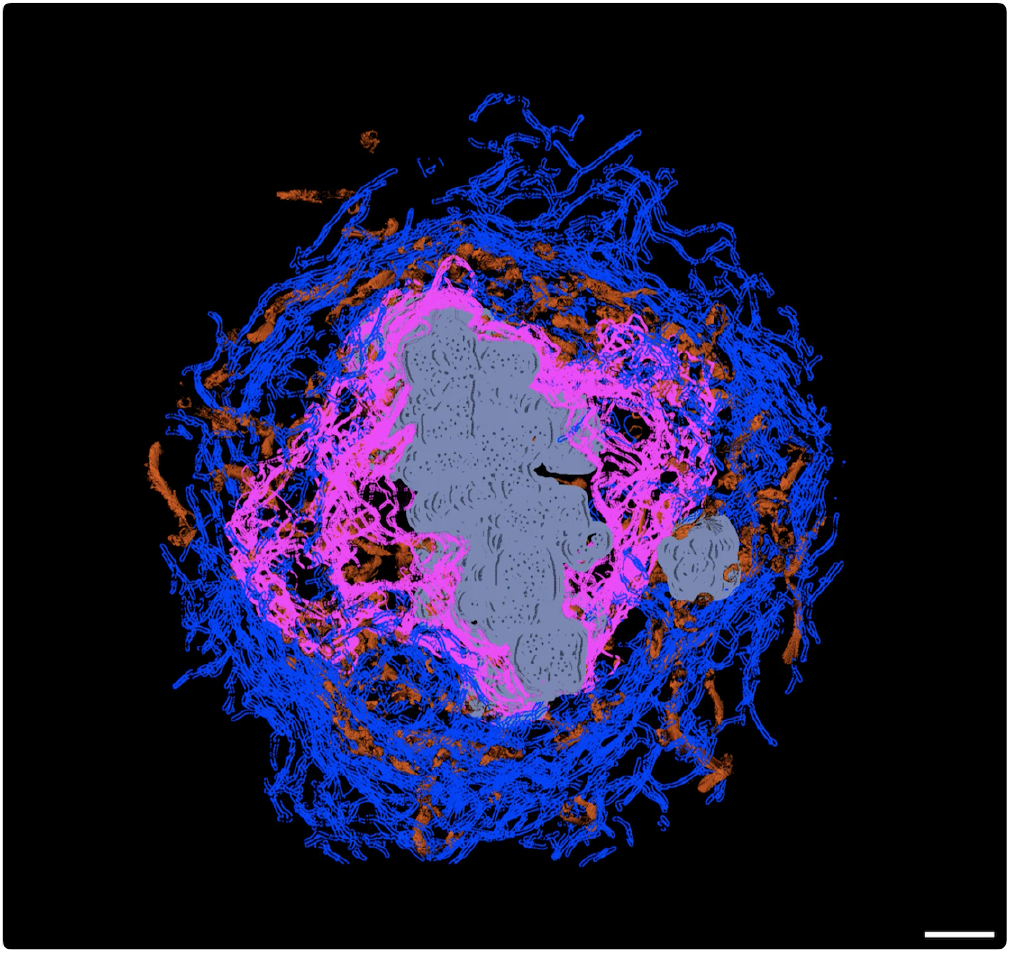
3D reconstruction of an ensheathed chromosome in an RPE-1 cell. A substack from SBF-SEM imaging showing a chromosome (gray) outside the exclusion zone (pink), ensheathed in endomembranes (blue). Three complete rotations are shown with DNA only, DNA plus exclusion zone boundary, finally with endomembranes (ER and mitochrondria, brown) added. Scale bar, 2 µm.

**Figure SV4.**
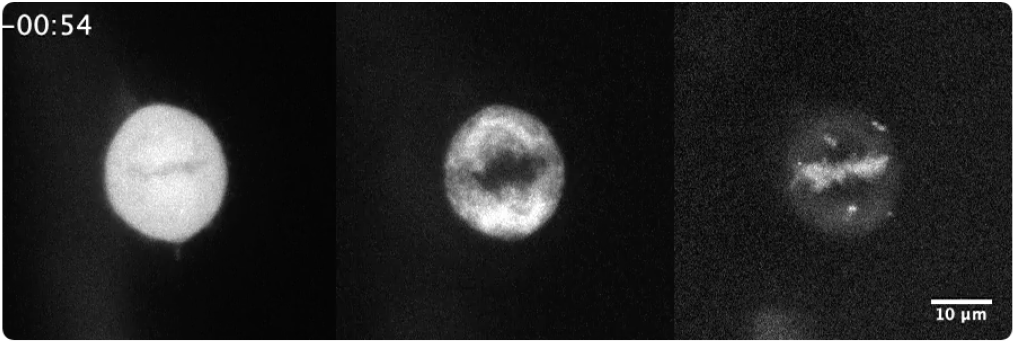
Example of GFP-Mad2 at an ensheathed chromosome. Movie of a GSK923295-pretreated RPE-1 cell stably expressing GFP-Mad2 (left) and mCherry-Sec61β (middle) with DNA stained with SiR-DNA (right). Time, hh:mm.

**Figure SV5.**
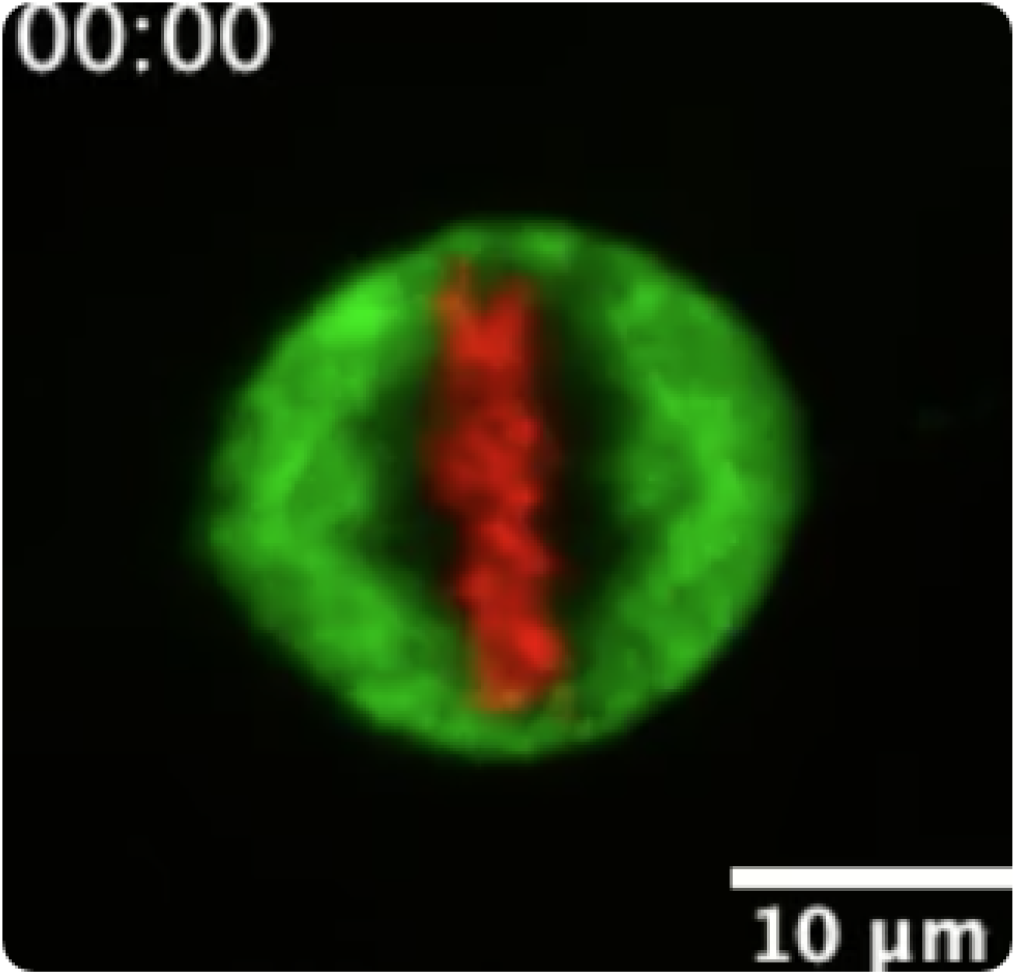
Example of mitotic outcome of a cell with aligned chromosomes. Movie of a control RPE-1 cell expressing GFP-Sec61β (green) stained with SiR-DNA (red). Cell has all chromosomes aligned and divides normally. Time, hh:mm.

**Figure SV6.**
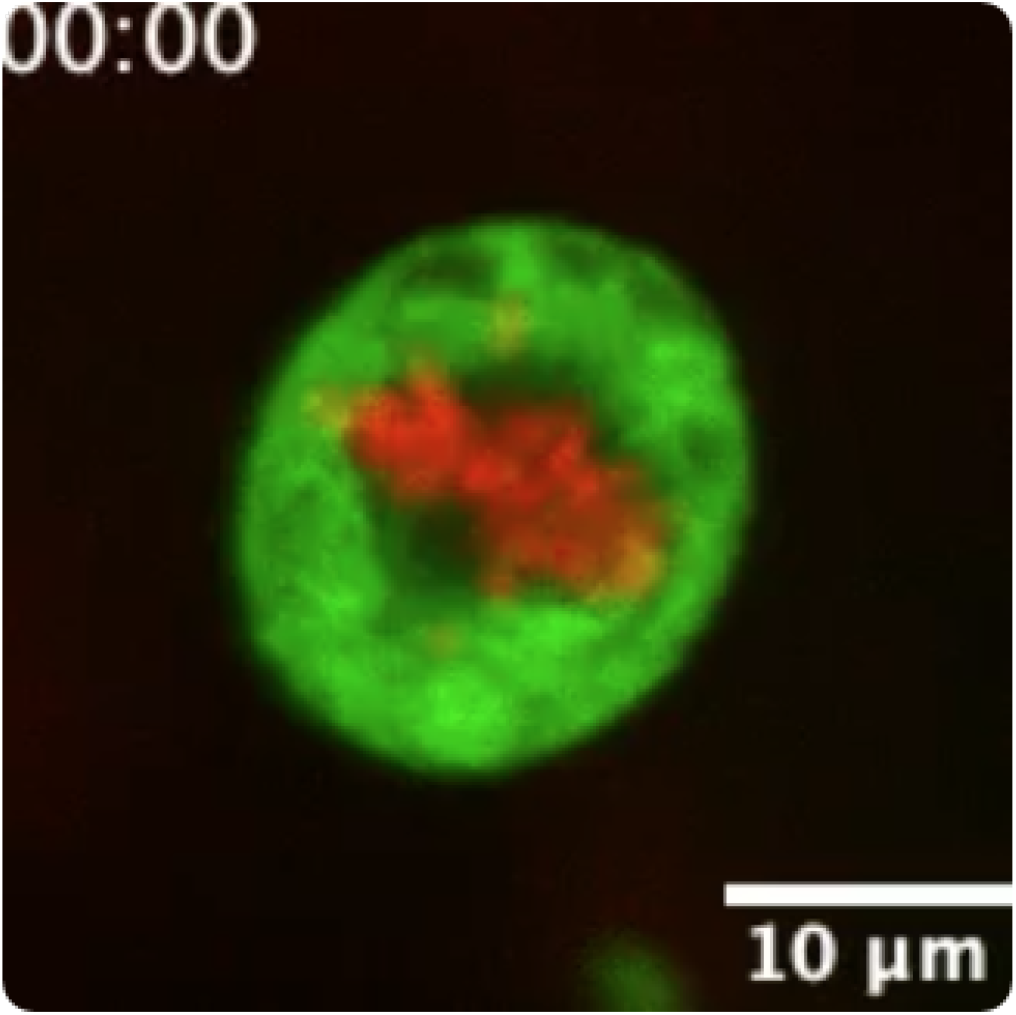
Example of mitotic outcome of a cell with an ensheathed chromosome. Movie of GSK923295 pre-treated RPE-1 cell expressing GFP-Sec61β (green) stained with SiR-DNA (red). Cell has an ensheathed chromosome and missegregates, leading to a micronucleus. Time, hh:mm.

**Figure SV7.**
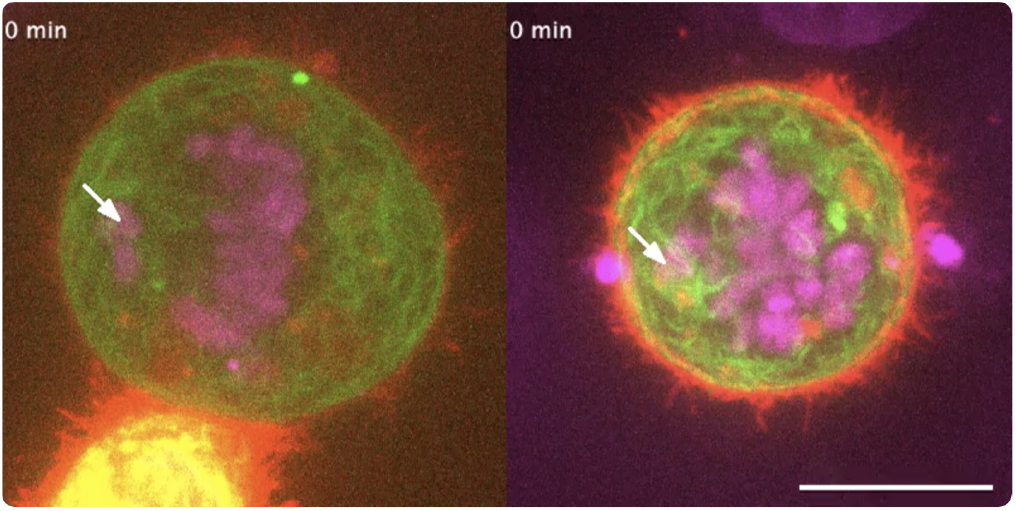
Example of ER clearance and subsequent rescue of an ensheathed chromosome. Movies of control (left) and ER clearance (right) in mitotic HCT116 cells expressing FKBP-GFP-Sec61β (green) and Stargazin-mCherry-FRB (red). DNA is stained with SiR-DNA (magenta). Scale bar, 10 µm.

## Notes

https://github.com/quantixed/Misseg

### Competing Interest Statement

The authors have declared no competing interest.

### Summary of Updates

Analysis of Mad2 levels of misaligned chromosomes. EM images to show ER relocalization to plasma membrane. Supplementary Information and movies updated.

## Bibliography

Bajer, A. (1957). Ciné-micrographic studies on mitosis in endosperm. III. The origin of the mitotic spindle. Exp Cell Res 13, 493–502. doi: 10.1016/0014-4827(57)90078-2.

Bakhoum, S. F., Ngo, B., Laughney, A. M., Cavallo, J.-A., Murphy, C. J., Ly, P., Shah, P., Sriram, R. K., Watkins, T. B. K., Taunk, N. K., Duran, M., Pauli, C., Shaw, C., Chadalavada, K., Rajasekhar, V. K., Genovese, G., Venkatesan, S., Birkbak, N. J., McGranahan, N., Lundquist, M., LaPlant, Q., Healey, J. H., Elemento, O., Chung, C. H., Lee, N. Y., Imielenski, M., Nanjangud, G., Pe’er, D., Cleveland, D. W., Powell, S. N., Lammerding, J., Swanton, C., and Cantley, L. C. (2018). Chromosomal instability drives metastasis through a cytosolic DNA response. Nature 553, 467–472. doi: 10.1038/nature25432.

Belevich, I., Joensuu, M., Kumar, D., Vihinen, H., and Jokitalo, E. (2016). Microscopy Image Browser: A Platform for Segmentation and Analysis of Multidimensional Datasets. PLoS Biol 14, e1002340. doi: 10.1371/journal.pbio.1002340.

Champion, L., Linder, M. I., and Kutay, U. (2017). Cellular Reorganization during Mitotic Entry. Trends Cell Biol 27, 26–41. doi: 10.1016/j.tcb.2016.07.004.

Champion, L., Pawar, S., Luithle, N., Ungricht, R., and Kutay, U. (2019). Dissociation of membrane-chromatin contacts is required for proper chromosome segregation in mitosis. Mol Biol Cell 30, 427–440. doi: 10.1091/mbc.E18-10-0609.

Cheeseman, L. P., Harry, E. F., McAinsh, A. D., Prior, I. A., and Royle, S. J. (2013). Specific removal of TACC3-ch-TOG-clathrin at metaphase deregulates kinetochore fiber tension. J Cell Sci 126, 2102–2113. doi: 10.1242/jcs.124834.

Cimini, D., Howell, B., Maddox, P., Khodjakov, A., Degrassi, F., and Salmon, E. D. (2001). Merotelic kinetochore orientation is a major mechanism of aneuploidy in mitotic mammalian tissue cells. J Cell Biol 153, 517–527. doi: 10.1083/jcb.153.3.517.

Clarke, N. I. and Royle, S. J. (2018). FerriTag is a new genetically-encoded inducible tag for correlative light-electron microscopy. Nat Commun 9, 2604. doi: 10.1038/s41467-018-04993-0.

Crasta, K., Ganem, N. J., Dagher, R., Lantermann, A. B., Ivanova, E. V., Pan, Y., Nezi, L., Protopopov, A., Chowdhury, D., and Pellman, D. (2012). DNA breaks and chromosome pulverization from errors in mitosis. Nature 482, 53–58. doi: 10.1038/nature10802.

Daum, J. R., Potapova, T. A., Sivakumar, S., Daniel, J. J., Flynn, J. N., Rankin, S., and Gorbsky, G. J. (2011). Cohesion fatigue induces chromatid separation in cells delayed at metaphase. Curr Biol 21, 1018–1024. doi: 10.1016/j.cub.2011.05.032.

Duijf, P. H. G. and Benezra, R. (2013). The cancer biology of whole-chromosome instability. Oncogene 32, 4727–4736. doi: 10.1038/onc.2012.616.

Dunlop, M. H., Ernst, A. M., Schroeder, L. K., Toomre, D. K., Lavieu, G., and Rothman, J. E. (2017). Land-locked mammalian Golgi reveals cargo transport between stable cisternae. Nat Commun 8, 432. doi: 10.1038/s41467-017-00570-z.

Dunsch, A. K., Linnane, E., Barr, F. A., and Gruneberg, U. (2011). The astrin-kinastrin/SKAP complex localizes to microtubule plus ends and facilitates chromosome alignment. J Cell Biol 192, 959–968. doi: 10.1083/jcb.201008023.

Fujiwara, T., Bandi, M., Nitta, M., Ivanova, E. V., Bronson, R. T., and Pellman, D. (2005). Cytokinesis failure generating tetraploids promotes tumorigenesis in p53-null cells. Nature 437, 1043–1047. doi: 10.1038/nature04217.

Funk, L. C., Zasadil, L. M., and Weaver, B. A. (2016). Living in CIN: Mitotic Infidelity and Its Consequences for Tumor Promotion and Suppression. Dev Cell 39, 638–652. doi: 10.1016/j.devcel.2016.10.023.

Ghadimi, B. M., Sackett, D. L., Difilippantonio, M. J., Schröck, E., Neumann, T., Jauho, A., Auer, G., and Ried, T. (2000). Centrosome amplification and instability occurs exclusively in aneuploid, but not in diploid colorectal cancer cell lines, and correlates with numerical chromosomal aberrations. Genes Chromosomes Cancer 27, 183–190.

Hatch, E. M., Fischer, A. H., Deerinck, T. J., and Hetzer, M. W. (2013). Catastrophic nuclear envelope collapse in cancer cell micronuclei. Cell 154, 47–60. doi: 10.1016/j.cell.2013.06.007.

Hepler, P. K. and Wolniak, S. M. (1984). Membranes in the mitotic apparatus: their structure and function. Int Rev Cytol 90, 169–238. doi: 10.1016/s0074-7696(08)61490-4.

Hirst, J., Edgar, J. R., Borner, G. H. H., Li, S., Sahlender, D. A., Antrobus, R., and Robinson, M. S. (2015). Contributions of epsinR and gadkin to clathrin-mediated intracellular trafficking. Mol Biol Cell 26, 3085–3103. doi: 10.1091/mbc.E15-04-0245.

Kalitsis, P., Earle, E., Fowler, K. J., and Choo, K. H. (2000). Bub3 gene disruption in mice reveals essential mitotic spindle checkpoint function during early embryogenesis. Genes Dev 14, 2277–2282. doi: 10.1101/gad.827500.

Kremer, J. R., Mastronarde, D. N., and McIntosh, J. R. (1996). Computer visualization of three-dimensional image data using IMOD. J Struct Biol 116, 71–76. doi: 10.1006/jsbi.1996.0013.

Kumar, D., Golchoubian, B., Belevich, I., Jokitalo, E., and Schlaitz, A.-L. (2019). REEP3 and REEP4 determine the tubular morphology of the endoplasmic reticulum during mitosis. Mol Biol Cell 30, 1377–1389. doi: 10.1091/mbc.E18-11-0698.

Larocque, G., La-Borde, P. J., Clarke, N. I., Carter, N. J., and Royle, S. J. (2020). Tumor protein D54 defines a new class of intracellular transport vesicles. J Cell Biol 219. doi: 10.1083/jcb.201812044.

Liu, S., Kwon, M., Mannino, M., Yang, N., Renda, F., Khodjakov, A., and Pellman, D. (2018). Nuclear envelope assembly defects link mitotic errors to chromothripsis. Nature 561, 551–555. doi: 10.1038/s41586-018-0534-z.

Lu, L., Ladinsky, M. S., and Kirchhausen, T. (2009). Cisternal organization of the endoplasmic reticulum during mitosis. Mol Biol Cell 20, 3471–3480. doi: 10.1091/mbc.e09-04-0327.

Lu, L., Ladinsky, M. S., and Kirchhausen, T. (2011). Formation of the postmitotic nuclear envelope from extended ER cisternae precedes nuclear pore assembly. J Cell Biol 194, 425–440. doi: 10.1083/jcb.201012063.

Ly, P., Teitz, L. S., Kim, D. H., Shoshani, O., Skaletsky, H., Fachinetti, D., Page, D. C., and Cleveland, D. W. (2017). Selective Y centromere inactivation triggers chromosome shattering in micronuclei and repair by non-homologous end joining. Nat Cell Biol 19, 68–75. doi: 10.1038/ncb3450.

Mammel, A. E., Huang, H. Z., Gunn, A. L., Choo, E., and Hatch, E. M. (2021). Chromo-some length and gene density contribute to micronuclear membrane stability. preprint, Cell Biology. URL http://biorxiv.org/lookup/doi/10.1101/2021.05.12.443914.

Merta, H., Carrasquillo Rodríguez, J. W., Anjur-Dietrich, M. I., Granade, M. E., Vitale, T., Harris, T. E., Needleman, D. J., and Bahmanyar, S. (2021). A CTDNEP1-lipin 1-mTOR regulatory network restricts ER membrane biogenesis to enable chromosome motions necessary for mitotic fidelity. preprint, Cell Biology. URL http://biorxiv.org/lookup/doi/10.1101/2021.03.02.433553.

Nicholson, J. M. and Cimini, D. (2015). Link between aneuploidy and chromosome instability. Int Rev Cell Mol Biol 315, 299–317. doi: 10.1016/bs.ircmb.2014.11.002.

Nixon, F. M., Honnor, T. R., Clarke, N. I., Starling, G. P., Beckett, A. J., Johansen, A. M., Brettschneider, J. A., Prior, I. A., and Royle, S. J. (2017). Microtubule organization within mitotic spindles revealed by serial block face scanning electron microscopy and image analysis. J Cell Sci 130, 1845–1855. doi: 10.1242/jcs.203877.

Porter, K. R. and Machado, R. D. (1960). Studies on the endoplasmic reticulum. IV. Its form and distribution during mitosis in cells of onion root tip. J Biophys Biochem Cytol 7, 167–180. doi: 10.1083/jcb.7.1.167.

Puhka, M., Vihinen, H., Joensuu, M., and Jokitalo, E. (2007). Endoplasmic reticulum remains continuous and undergoes sheet-to-tubule transformation during cell division in mammalian cells. J Cell Biol 179, 895–909. doi: 10.1083/jcb.200705112.

Puhka, M., Joensuu, M., Vihinen, H., Belevich, I., and Jokitalo, E. (2012). Progressive sheet-to-tubule transformation is a general mechanism for endoplasmic reticulum partitioning in dividing mammalian cells. Mol Biol Cell 23, 2424–2432. doi: 10.1091/mbc.E10-12-0950.

Schlaitz, A.-L., Thompson, J., Wong, C. C. L., Yates, J. R., and Heald, R. (2013). REEP3/4 ensure endoplasmic reticulum clearance from metaphase chromatin and proper nuclear envelope architecture. Dev Cell 26, 315–323. doi: 10.1016/j.devcel.2013.06.016.

Schweizer, N., Pawar, N., Weiss, M., and Maiato, H. (2015). An organelle-exclusion envelope assists mitosis and underlies distinct molecular crowding in the spindle region. J Cell Biol 210, 695–704. doi: 10.1083/jcb.201506107.

Uetake, Y. and Sluder, G. (2010). Prolonged prometaphase blocks daughter cell proliferation despite normal completion of mitosis. Curr Biol 20, 1666–1671. doi: 10.1016/j.cub.2010.08.018.

van Bergeijk, P., Adrian, M., Hoogenraad, C. C., and Kapitein, L. C. (2015). Optogenetic control of organelle transport and positioning. Nature 518, 111–114. doi: 10.1038/nature14128.

Vedrenne, C., Klopfenstein, D. R., and Hauri, H.-P. (2005). Phosphorylation controls CLIMP-63-mediated anchoring of the endoplasmic reticulum to microtubules. Mol Biol Cell 16, 1928–1937. doi: 10.1091/mbc.e04-07-0554.

Wang, X., Le, N., Denoth-Lippuner, A., Barral, Y., and Kroschewski, R. (2016). Asymmetric partitioning of transfected DNA during mammalian cell division. Proc Natl Acad Sci U S A 113, 7177–7182. doi: 10.1073/pnas.1606091113.

Warren, G. (1993). Membrane partitioning during cell division. Annu Rev Biochem 62, 323–348. doi: 10.1146/annurev.bi.62.070193.001543.

Wood, K. W., Lad, L., Luo, L., Qian, X., Knight, S. D., Nevins, N., Brejc, K., Sutton, D., Gilmartin, A. G., Chua, P. R., Desai, R., Schauer, S. P., McNulty, D. E., Annan, R. S., Belmont, L. D., Garcia, C., Lee, Y., Diamond, M. A., Faucette, L. F., Giardiniere, M., Zhang, S., Sun, C.-M., Vidal, J. D., Lichtsteiner, S., Cornwell, W. D., Greshock, J. D., Wooster, R. F., Finer, J. T., Copeland, R. A., Huang, P. S., Morgans, D. J., Dhanak, D., Bergnes, G., Sakowicz, R., and Jackson, J. R. (2010). Antitumor activity of an allosteric inhibitor of centromere-associated protein-E. Proc Natl Acad Sci U S A 107, 5839–5844. doi: 10.1073/pnas.0915068107.

Yang, Z., Loncarek, J., Khodjakov, A., and Rieder, C. L. (2008). Extra centrosomes and/or chromosomes prolong mitosis in human cells. Nat Cell Biol 10, 748–751. doi: 10.1038/ncb1738.

